# Spontaneous eye-movements during eyes-open rest reduce resting-state-network modularity by increasing visual-sensorimotor connectivity

**DOI:** 10.1101/2020.05.18.100669

**Authors:** Cemal Koba, Giuseppe Notaro, Sandra Tamm, Gustav Nilsonne, Uri Hasson

## Abstract

During wakeful rest, individuals make small eye movements when asked to fixate. We examined how these endogenously-driven oculomotor patterns impact topography and topology of functional brain networks. We used a dataset consisting of eyes-open resting-state (RS) fMRI data with simultaneous eye-tracking (Nilsonne et al., 2016). The eye-tracking data indicated minor movements during rest, on the order of 1.0 degree on average when analyzed over 2sec epochs, which correlated modestly with RS BOLD data. However, the eye-tracking data correlated well with echo-planar imaging (EPI) time series sampled from the area of the Eye-Orbit (EO-EPI), which is a signal previously used to identify eye movements during exogenous saccades and movie viewing. We found that EO-EPI data correlated with activity in an extensive motor and sensory-motor network, but also some components of the dorsal attention network including the frontal and supplementary eye fields. Partialling out variance related to EO-EPI from RS data reduced connectivity, primarily between sensory-motor and visual areas. For three different network sparsity levels, the resulting RS connectivity networks showed higher modularity, lower mean connectivity strength, and lower mean clustering coefficient. Our results highlight new aspects of endogenous eye movement control during wakeful rest. They show that oculomotor-related contributions form an important component of RS network topology, and that those should be considered in interpreting differences in network structure between populations, or as a function of different experimental conditions.

## Introduction

The study of human brain activity during resting state (RS) is of considerable interest in both basic and clinical brain research. For mechanistically-oriented perspectives, RS activity patterns identify constraints that may govern task-evoked activity as seen by relations between RS connectivity and inter-individual differences in various cognitive tasks (e.g., Kelly, Uddin, Biswal, Castellanos, & Milham, 2008; Rosenberg, Hsu, Scheinost, Constable, & Chun, 2018). And because RS connectivity is related to structural connectivity (Honey et al., 2009; Mišić et al., 2016), it is considered an important mediator between anatomical organization and task-evoked activity. From the perspective of predictive models of interindividual differences in healthy and clinical populations, the quantification of RS features (using time-domain, network-based analyses, spatiotemporal clustering, control-based approaches, to name a few) is used for machine-learning or statistical learning. This has proved promising in contexts such as prediction of IQ (Dubois, Galdi, Paul, & Adolphs, 2018), personality (e.g., Nostro et al., 2018), or the likelihood of developing clinical conditions (e.g., de Vos et al., 2018).

RS data measured via fMRI reflect endogenous neural activity, but also additional sources such as physiological artifacts (e.g., cardiac and respiratory effects, Birn, 2012; J. Chen et al., 2020), or head and body motion (e.g., Parkes, Fulcher, Yücel, & Fornito, 2018). For machine learning, these non-neural effects on the BOLD signal may be informative (e.g., motion-related patterns could differ across populations (e.g., Zacà, Hasson, Minati, & Jovicich, 2018). However, motion and physiological effects complicate drawing conclusions about brain systems mediating endogenous information-computation during wakeful rest. For this reason, researchers often remove effects of motion and physiology from RS data prior to analysis.

Here we examined how RS connectivity is related to a different factor, which is eye movement during rest. For purposes of understanding endogenous computations, spontaneous eye-movement at rest straddles the space between an interesting neurobiological phenomenon reflecting the output of endogenous activity and a nuisance factor reflecting motor activity. On one hand, eye-movement can be considered a truly integral component of wakeful rest, because at minimum, retinal input is continuously refreshed to minimize adaptation. On the other hand, oculomotor control differs from prototypical covert, non-motor processes exactly because motor control involves planning, execution, efference copy, feedback and correction (e.g., West, Welsh, & Pratt, 2009). Oculomotor-control during rest may therefore instantiate coordination between brain systems that otherwise present modest levels of coordination.

Statistically, eye movements during rest could produce stronger connectivity between regions. Perhaps more importantly, it could produce a more integrated (less-modular) view of RS connectivity networks, because eye movement is supported by a widely distributed fronto-parietal network and occipital regions (e.g., Balslev, Albert, & Miall, 2011; Mort et al., 2003). From a theoretical perspective, identifying neural systems controlling eye movement during rest could allow better partitioning between relatively more ‘active’, (oculo)motor-related aspect of RS as opposed to other more covert, non-motor-related aspects of RS. Finally, eye-movement themselves could be a possible confounder when studying controls and clinical groups that differ in oculomotor control including Autism (e.g., Takarae, Minshew, Luna, Krisky, & Sweeney, 2004) or Parkinson’s Disease (e.g., Pretegiani & Optican, 2017; Zhang et al., 2018).

### Current knowledge

There is relatively little prior work on the relationship between eye movements and RS activity. Using fMRI, Fransson, Flodin, Seimyr, and Pansell (2014) studied neural correlates of horizontal or vertical guided fixations, as well as spontaneous fixations during RS. Guided fixations produced activity in systems typically involved in oculomotor movement including visual cortex, frontal eye fields (FEF), supplementary motor area (SMA), cerebellum, and a few other regions. To quantify correlates of spontaneous eye movement during RS they derived a gaze-velocity time series from the eye tracking data, reduced its dimensionality using PCA, convolved the resulting timeseries with a hemodynamic response function and used the result as a regressor in a whole-brain analysis. Interestingly, this latter analysis identified fewer regions that did not overlap with those found for guided saccades, and which were all associated with the Default Mode Network (DMN): the posterior cingulate cortex (PCC) and dorsomedial prefrontal cortex (dmPFC). As the authors noted (p. 3833), “at first glance it would seem more likely to expect the neuronal control for slow changes in eye position during fixation to be localized to visual cortices and attention-related cortical networks”. It it unclear how slow fluctuations in the DMN impact eye movement.

McAvoy et al. (2012) used Electro-oculography (EOG) to monitor eye movement during fixation, in an analysis based on a relatively small sample (*N* = 9). Using the EOG they separated blinks from other eye movement during eyes-open RS. In the analysis of EOG during RS fixation they identified brain systems correlated with blinks, but no brain systems where activity correlated with other types of eye movements.

Yellin, Berkovich-Ohana, and Malach (2015) examined correlations between blood oxygen level-dependent (BOLD) fluctuations during rest and pupil size. They identified widespread negative correlations in sensory-motor areas and temporal areas, and positive correlations in the DMN. The study did not evaluate BOLD correlates of gaze location or velocity. However, it is possibly related to understanding systems related to spontaneous eye movement, because pupil-size measurements are known to be confounded with the deviation of the pupil from the center of camera view. That is, eye trackers will mis-report systematically decreasing pupil-size values – for the exact same pupil size – as a function of the deviation of pupil from camera-axis (Hayes & Petrov, 2016).^1^

Ramot et al. (2011) used EOG to determine BOLD correlates of spontaneous eye movements during an eyes-closed condition. the relation to eyes-open oculomotor control is unclear, as eyes-closed RS conditions produce different activity (e.g., Marx et al., 2003) and connectivity (e.g., McAvoy et al., 2012) patterns. Furthermore, saccades made under closed eye lids have different trajectories than those made with eyes open in complete darkness (Becker & Fuchs, 1969). For this reason we consider prior studies examining RS activity during eyes-open condition as more relevant for current work.

In addition, numerous neuroimaging studies have used various types of tasks, including visually-guided saccades, memory-guided saccades, anti-saccades and so-called “voluntary” saccades (either pre-cued [endogenous control] or freely initiated). However these studies used explicit tasks rather than study naturally occurring oculomotor control during eyes-open RS. Perhaps the essential difference is that controlled studies oftentimes orient the saccade towards, or away from a presented target (pro-vs anti-saccade) and for this reason the brain systems identified could mediate visual detection and attention processes that have no parallel during rest. In a neuroimaging study demonstrating this point, the authors (Brown, Goltz, Vilis, Ford, & Everling, 2006) required participants to saccade either towards a stimulus (prosaccade), away from a stimulus (antisaccade), or maintain fixation while inhibiting an orienting saccade (no-go). They documented numerous regions, including FEF, IPS, cingulate cortex and precuneus, all showing highly similar activation patterns for both prosaccade and no-go trials. The authors wrote this suggests that “BOLD signal in cortical saccade regions might predominantly reflect visual detection and attention processes rather than saccade generation or inhibition… “For this reason, it is unclear to what extent brain systems identified in typical studies of saccades are strongly involved in saccade control during the resting state.

### Specific aims

The two aims of our current study were: 1) to identify brain systems associated with endogenously driven eye movements during rest, and conjointly, 2) to determine how removal of eye-movement related activity impacts resting-state connectivity. We quantified eye movement during rest using both eye-tracking, and EPI data extracted from the eye orbit area. We validated the relationship between different features of oculomotor movement (pupil size, gaze velocity, gaze location) and Eye Orbit EPI time series (EO-EPI) during rest. We then evaluated how removal of eye-related activity, as manifested in EO-EPI, impacts the topography and topology of RS networks.

## Methods

### Dataset

We used resting state data from the Sleepy Brain study (Nilsonne et al., 2016).^2^ Full details of the dataset and imaging parameters are given in Nilsonne et al. (2016) and here we provide only the main details. Data were collected from 86 participants on a 3T MRI scanner (Discovery 750, General Electric) using an 8-channel head coil. Each participant was scanned on two different days. In each scanning session, a T1 structural image, two resting state functional EPI scans, and three task-related functional scans (emotional mimicry, empathy for pain, emotional reappraisal) were acquired. Our analyses rely only on the structural and resting-state scans.

For the structural (T1) images, the relevant properties were as follows: slice thickness 1mm, sagittal orientation, whole brain acquisition. For the resting state EPI images: slice thickness 3mm no gap, axial orientation, 49 slices covering the entire brain, interleaved acquisition inferior to superior, *TE* = 30, *TR* = 2.5*s*, flip angle 75*°*.

Four resting-state data sets were acquired for each participant; two runs on each of two scanning days. In one of the two days, data were collected when participants were sleep deprived, and we did not analyze these data. Of the remaining two RS runs, one was typical, where participants were asked to fixate on a white cross presented a gray background for 8 minutes. The second run was quasi-rest in that in addition to fixation, it included self-rated sleepiness probes every two minutes. We only analyzed data from the typical RS session. To summarize, we processed one RS run per participant, which was a typical RS scan acquired in absence of sleep deprivation. Three participants did not provide these runs so 83 participants were included in our initial sample. Participants belonged to two age groups: 20–30 and 65–75 y.o.a.

### Pre-processing of eye tracking data

Eye tracking data were available for 77 of the 83 participants for which we analyzed the RS data. Participants were required to maintain their gaze on a central fixation cross for the duration of the 8 min scan. Right eye movement and pupil size were recorded using an Arrington Research Viewpoint system integrated into head-mounted goggles. Eye data were sampled at 60 Hz. Participants were monitored during the experiment to ensure that they did not have prolonged eye closures (*>* 5s).

When analyzing these data we observed a substantial proportion of missing values, likely due to loss of pupil tracking during the task. We therefore implemented a quality assurance procedure as detailed below. As recommended by the instrument manufacturer, we detected eyeblinks by identifying local deviations (anomalies) in the values of pupil sizes. Specifically, we defined blink artifacts as cases where *i*) the ratio of the pupil width to pupil height (pupil aspect) was too excessive, or when one of the pupil dimensions exceeded the validity range that, by visual inspection, was in the range 0.1-0.5 (instrumental arbitrary units). To avoid to reliance on arbitrary thresholds, we defined an artifact function as the sum of the following three functions (each normalized to its maximum value). In these equations, *f*_1_ is the pupil aspect ratio, and *f*_2_ and *f*_3_ diverge when one pupil dimension approach the boundaries of the validity range 0.1-0.5.

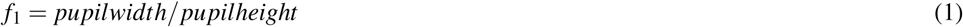

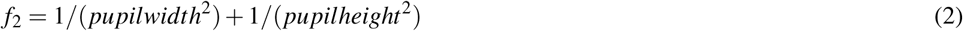

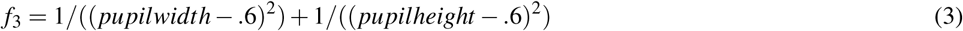

After transforming the using these functions, we defined artifacts as sharp peaks in the resulting time series. In addition, we defined blinks as artifacts whose duration was 100–400 msec. After removing time points containing artifacts, we considered the time series of gaze locations. To limit the influence of the noise due to data acquisition failures we only included data from participants with gaze variance below an arbitrary value of 0.1 radians. Consequently, we analyzed data from 32 (of 77) participants. For these, the proportion of artifacts was on average 18 *±* 2%; blinks occurred with an average period of 2.36 *±* 0.21 sec.

### Pre-processing of fMRI data and creation of eye-orbit EPI regressors

We include the analysis workflow described below as supplementary materials, also available online.

First, we applied brain extraction and tissue segmentation (Gray Matter, White Matter, CSF) to the structural T1 images using the *antsBrainExtraction* function of ANTs software (Avants, Tustison, & Song, 2011). We used ANTs for all registration routines in our pipeline. We registered each participant’s structural image to standard space using non-linear registration (ICBM 2009 non-linear assymetric template; Fonov, Evans, McKinstry, Almli, & Collins, 2009), and saved the inverse of the warps. We also registered the structural and functional images using affine transformation. We used the combination of these two transformations to align data from each participant’s original space to common space, or vice versa, in a single step.

To delineate each participants “eye orbit” area we first marked this area on the common-space template. We then transformed this mask to each participant’s original space, and made any additional modifications (if needed) therein. Specifically, we delineated anatomical masks of the “eye orbit” area in common space using MRICRON (Rorden, Karnath, & Bonilha, 2007), for which we used an MNI template provided with FSL (Jenkinson, Beckmann, Behrens, Woolrich, & Smith, 2012). Both eye orbits were included in the mask. The masks’ location was transformed to each participant’s individual space using the combination of T1 → subject space matrices and inverse of the T1 → MNI matrices mentioned above. We also created cerebral-spinal fluid (CSF) and white matter masks in MNI space and transformed them to individual space, where they were eroded by one voxel from each direction to be more conservative. We then extracted the mean time series from these white matter and CSF masks. These were used as nuisance regressors in an initial regression (details below).

We used AFNI (Cox, 1996) for analyzing the functional RS images. We implemented the following steps: slice time correction, motion correction (base image: first volume of the run), and band-pass filtering (0.01 − 0.1*Hz*). To remove other nuisance sources of variance from the functional times series we implemented preliminary data-cleaning using regression with the following regressors: *i*) motion parameters estimated during motion correction, *ii*) mean white matter and CSF time series, *iii*) and frame-wise displacement values (included in the model as a regressor). We considered the residuals of this regression as a “cleaned” time series that was the starting point for further analyses.

To improve signal to noise of the subsequent regression models which were of primary interest, we then spatially-smoothed the cleaned time series with a 6mm FWHM kernel. From this time series we also derived an Eye-Orbit EPI regressor, which was defined as the mean time series from both eye-orbit masks, after spatial smoothing, which we refer to as EYE_*raw*_. We convolved the EYE_*raw*_ with an HRF basis function (Using AFNI’s *waver* command), producing a EYE_*conv*_ time series. In separate analyses we used either EYE_*raw*_ or EYE_*conv*_ as “seed” regressors, to identify EO-EPI-correlated brain areas.

### Determining correlation between eye-tracking measures and EO-EPI time series

We were interested in the relationship between several measures of eye movement and the EPI time series sampled from the eye-orbit regions (EO-EPI series). We derived 12 time series from the eye-tracking data were: the measured gaze location, *GazeX* and *GazeY* (mean normalized for horizontal center per participant), their squared values, their temporal derivatives (*vel GazeX, vel GazeY*), gaze amplitude: *GazeX* ^2^ + *GazeY* ^2^, gaze power: *vel GazeX* ^2^ + *vel GazeY* ^2^, *Pupil size* (de-meaned), its first derivative *vel Pupil size*, and squared value *Pupil size*^2^. We were also interested in the *blink function* (coding for 1 whenever a blink was present; 0 otherwise), but we determined the relation between blinks and EO-EPI in a different manner as detailed below. ^3^

For each of the 12 eye-tracking quantities mentioned above (with the exception of blinks) we performed the following procedure: We first down-sampled the time series to the fMRI frequency rate (0.4 Hz). Rather than a-priori assume that the relation between the eye-tracker data and EO-EPI is mediated by a typical hemodynamic response function, we used a simple statistical learning approach to estimate and validate this relationship. Specifically, we calculated a kernel function to describe the relation between the eye tracking quantity and the EO-EPI envelope. To calculate a kernel, we implemented the following procedure. First, for each oculomotor time series we considered as meaningful oculomotor ‘events’ the top 10% of the peak-values in the given series. Second, we calculated the mean EO-EPI signal in the interval [−10, 10] seconds around those peak events. For each participant, the triggered mean was normalized to that participant’s absolute maximum value, in this way producing the participant’s event triggered average (ETA). Finally, for each participant, we used the mean of the ETAs calculated from *all other* participants as the kernel to apply to the left-out participant’s data (this maintained independence of estimation and testing). The kernel was convolved with the eye tracking time series, and a correlation with EO-EPI computed. The resulting correlation values (32 in all) were then Fisher-Z transformed and analyzed on the group level using a T-test.

We used a different approach to evaluate the relation between blinks and EO-EPI dynamics. The blink time series was sparse and binary, with ‘1’ coding blink presence. We down-sampled this time series to consecutive 2.5 sec windows, assigning to each window the value 1 if at least one blink was coded in the original series. For each participant we computed a blink-related event-triggered-average by averaging the EO-EPI data around each blink (as described above). To determine the statistical significance of blinks and EO-EPI we evaluated the reliability of the ETAs across participants: We calculated for each participant the correlation between his/or own ETA and the average of the ETAs of all the other subjects. We then tested the distribution of these correlation values at the group level using a T-test.

### Statistical Inference for fMRI analyses

#### Correlates of Eye-tracking metrics

We examined whole-brain RS correlations with several eye tracking measures: *GazeX, GazeX* ^2^, *vel GazeX, vel GazeX* ^2^, *Pupil size*, and *blink function*. The BOLD data modeled were “cleaned” time series from which only typical artifact sources were removed. We implemented two modeling approaches: In the first, we resampled each eye-tracking measure of interest to the sampling resolution of the MR acquisition (*Hz* = 0.4) and convolved the result with canonical HRF via AFNI’s Waver function to construct a regressor. In the second, we used a Finite Impulse Response (FIR) function modeling approach where the BOLD impulse response was estimated using six tent functions (using AFNI’s *tent* basis function). This approach does not assume a fixed shape. From these estimates, we averaged the first three beta coefficients (corresponding to 0 − 7.5*sec* post eye-tracker dynamics) and propagated the value to a group-level analysis. Family wise error correction was implemented using FSL’s TFCE implementation.

#### Correlates of EO-EPI Regressors

Beta values associated with EYE_*conv*_ or EYE_*raw*_ were transformed to MNI space. To identify clusters where these beta values were significantly positive or significantly negative we computed voxel-wise statistics (Wilcoxon signed-rank test) and then implemented cluster-level control for family-wise-error using permtuations as described below. We used a non-parametric test because the relevant Beta values data did not satisfy typical parametric assumptions.

We defined statistically-significant clusters as ones where the statistical significance (uncorrected) at the single voxel level was below *p* = .01, and where the cluster size (volume) passed a value determined from the sampling distribution we derived using the following permutation procedure. In each of 10,000 permutations, we reversed the signs of 42 of the 83 datasets (i.e., the labels for conditions EYE_*conv*_ and EYE_*raw*_ were flipped for these participants), and we implemented a Wilcoxon signed-rank test (Siegel & Castellan, 1956) to identify all clusters consisting of voxels where the statistical significance of the difference from chance (zero; 0) exceeded *p <* .01 and where all values were positive (we limited to positive values so that the resulting clusters could not combine both negative and positive values, as our main analysis also probed for clusters where all values were either positive or negative). We saved the largest cluster size from each permutation, and the resulting set of 10,000 values of largest-cluster sizes defined the sampling distribution. The 95% percentile rank entry of the sampling distribution served as the critical value. This value was used to define statistically-significant clusters in the experimental data.

To evaluate whether significant EO-EPI correlates were found in areas dominated by artifacts, we calculated temporal signal to noise ratio (tSNR) for each participant. To create tSNR map for each participant, we used the raw functional images (before applying any signal processing steps), but after removal of the initial 10 stabiliziation images. We divided the absolute mean value of each voxel by its standard deviation. We then applied the statistically significant clusters found for EYE_*raw*_ and EYE_*conv*_ series as masks to determine *mean* of the tSNR in each statistically significant spatial cluster. The motivation for this analysis was a prior report (W. Chen & Zhu, 1997) showing that Nyquist ghosting artifacts can propagate eye signals into midbrain areas (in the case of axial acquisition). Two MR physicists examined the QA reports produced by the scanner and did not find evidence for ghosting. However, we still wanted to evaluate if any EO-EPI whole-brain correlates were found in regions with low tSNR.

We defined the frontal eye fields (FEF) and supplementary eye fields (SEF) as independent Regions of Interest and for each each we examined correlations with the EO-EPI regressor. To create FEF and SEF regions of interest, we used the NeuroSynth database (Yarkoni, Poldrack, Nichols, Van Essen, & Wager, 2011). The probability mask corresponding to the keyword *eye* was saved and thresholded by z-score of 7 (max Z=9.1, generated from 417 studies). From the thresholded image, regions around the intersection of precentral sulcus and superior frontal sulcus were marked as FEF, and region around medial frontal gyrus was marked as SEF (see Appx.4). Those masks were spatially translated to the individual-subject space and mean activation of those two ROIs extracted from the cleaned and smoothed data. We constructed a regression model to predict that FEF and SEF regional activity from the EO-EPI series per participant. Coefficients were analyzed using a Wilcoxon rank sum test.

#### Functional connectivity maps and derived network metrics

To create functional connectivity networks, we used Schaefer et al. (2018)’s resting-state functional connectivity parcellation based on 500 regions of interest (ROIs). We spatially translated this parcellation into each participant’s individual space, where they were further limited to gray matter by multiplying all ROIs with the participant-specific gray matter mask (to limit the influence of data from non-gray matter areas). We extracted the mean time series from each ROI, for the two types of spatially-smoothed resting-state data we derived (one typical, and the other with EO-EPI regressed). We examined the network features after thresholding the connectivity matrices at three sparsity levels – 10%, 20%, 30%. While the thresholding removed weak connections, we did not binarize the remaining (above-threshold) connections but analyzed the complete set of data. From each participant’s resting state network we derived the following metrics: node degree, strength, cluster coefficient, transitivity, assortativity, efficiency, number of communities, betweenness centrality and modularity. We calculated these using the Brain Connectivity Toolbox (Rubinov & Sporns, 2010) (See *Appendix* for description of the metrics as described in the Brain Connectivity Toolbox). We calculated these parameters for the original and “clean” networks as defined above. We then tested which of these parameters differed as a result of the EO-EPI-removal procedure using paired-sample T-tests. We defined a robust result as one that was statistically significant across all three levels of sparsity thresholding (full results are reported for completeness).

We also probed changes in global topology by quantifying the impact of EO-EPI removal on the shape of the entire degree distribution (within each sparsity level). Following prior work (e.g., Fornito, Zalesky, & Bullmore, 2010) we fit a truncated power law function to each participant’s degree distribution. The function was *Y* = *a × X*^*b*^ *× e*^(*x×c*)^, Where *Y* is the cumulative probability of the distribution and *x* = node degree. From this equation, we derived the coefficient (*a*), power law exponent (*b*), and degree cut-off point (*c*). A paired-sample t-test was applied for each parameter to see if the parameters differ across conditions.

Using previously defined criteria (Xu et al., 2014), we detected hubs for the networks defined by the three sparsity thresholds. These criteria required that the value of a node be higher than 1 SD above the mean value for each of these empirical distributions: node strength, node degree and node between-ness centrality. Nodes matching all three criteria were considered hubs. The chance probability of a node being a hub (assuming a normal distribution) is *∼*0.34^3^ = .04. To evaluate whether removal of EO-EPI variance impacted whether a region satisfied hub criteria, for each region we counted the number of participants for which the region was classified as a hub, with our without EO-EPI removal. On a binomial, a difference would need to consist of at least 7 or more participants (binomial test parameters: *N* = 86; *K* = 7; *p* = .04).

We also identified any specific pair-wise differences in regional connectivity for the raw and cleaned matrices. After applying Fisher’s Z transformation, pair-wise correlation values were subjected to paired-sample t-tests. We used false discovery rate (FDR) to correct for multiple comparisons.

#### Dual Regression

We used dual regression to determine how removal of activity associated with the EO-EPI regressor impacted connectivity in well-defined resting-state networks. The procedure was implemented in AFNI and followed workflows described previously (Beckmann, Mackay, Filippini, & Smith, 2009; Nickerson, Smith, Ö ngü r, & Beckmann, 2017). In the first step we used 14 pre-defined resting-state network spatial masks (Shirer, Ryali, Rykhlevskaia, Menon, & Greicius, 2012) to extract ‘seed’ time series for each of the networks. The 14 resting-state network masks were spatially transposed to individual space and multiplied with gray matter of the participant to reduce contribution from non-gray-matter areas. For each participant we then produced two seed time-series for each of the 14 networks: one from the functional data from which the EO-EPI variance was not removed, and one from the functional data from which this variance was removed using the EYE_*conv*_ regressor.

To determine whole brain connectivity of the seed regions we inserted all 14 time series into a single multiple regression. In effect, we conducted two separate regression models: Model #1 was a “typical” model where the mask-derived seed time series produced from the original (typically-processed) functional data served as regressors to predict whole-brain resting state data. This process replicates the procedure typically used in resting-state dual regression. Model #2 was an “EO-EPI-removed” model where masked-derived seed time series produced from EO-EPI-removed functional data were used as regressors to predict the EO-EPI-removed functional data. This is a dual regression that uses EO-EPI-cleaned resting state data.

The produced beta weights were analyzed using group level repeated-measures test to identify seed-time-series whose connectivity differed between the two data sets; i.e., whose connectivity was impacted by the EO-EPI-removal procedure. We used FSL’s *randomise* function (Jenkinson et al., 2012). A within group T-test with 10000 permutations and threshold-free cluster enhancement was applied. Because our interest was in evaluating the impact EO-EPI-regressor we adopted a liberal approach of not correcting for multiple comparisons across the 14 networks tested in the dual regression procedure. We also note that the 14 time series used for dual regression were relatively weakly correlated in this data set: to determine collinearity, on the single participant level we computed the 14 *×* 14 cross-correlation matrix and then averaged these across participants. The highest mean correlation was 0.55, which licensed separate analyses for each network regressor.

## Results

### Eye tracking data: Quality and correlation with whole-brain BOLD

Based on our artifact rejection criteria, usable eye-tracking data were available for 32 of 77 participants for which eye tracking data were collected. A power-spectra analysis of the eye tracking data (Figure Appx.1) indicated higher broad-band power in all frequencies in the rejected data, including those approaching the Nyquist frequency of the eye-tracking data in the current study (*f* = 30*Hz*). Participants largely avoided making large eye movements during the resting-state session. To quantify these movements, we calculated the maximal displacement of gaze position in non-overlapping 2sec windows. The resulting empirical cumulative distribution functions (see Figure 1A) indicated modest movement, with around 50% of analysis windows showing displacement values *<* 1*°* and only around 10% of windows showing displacement values *>* 3*°*.

**Figure 1.**
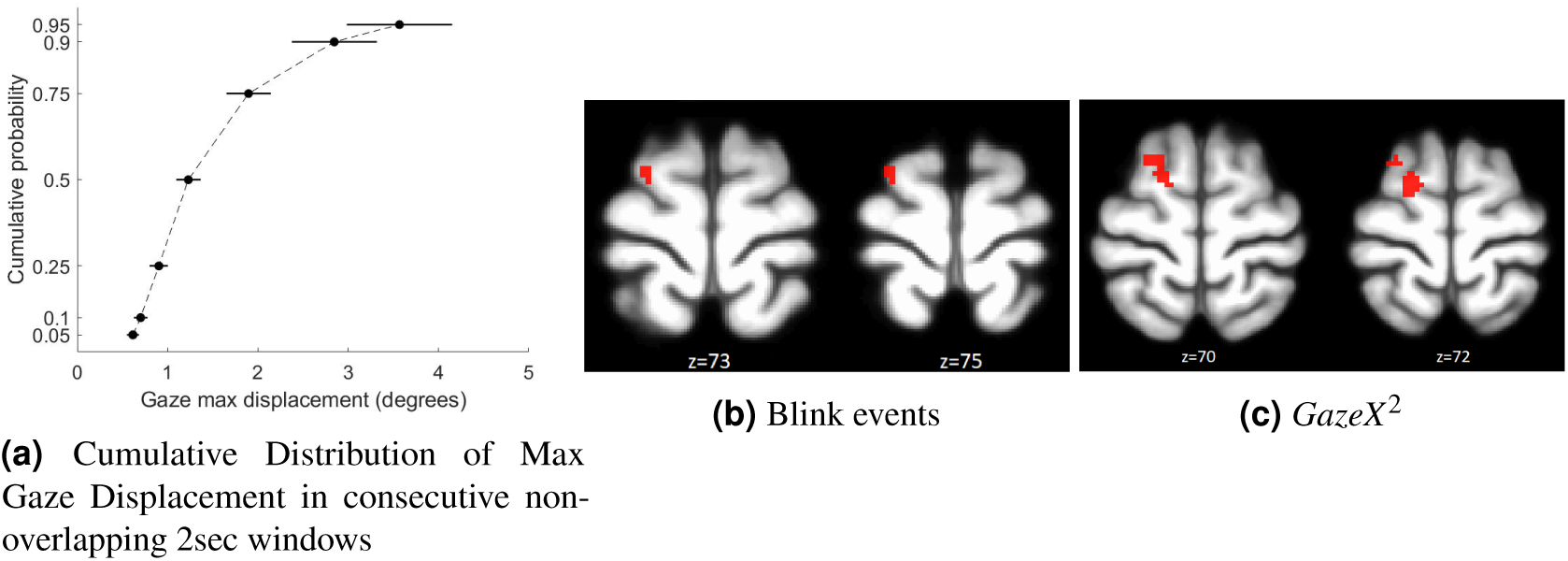
Relation between eye-tracking measures and EO-EPI regressor from eye orbits. Panel A: modest eye movements in 2-sec non-overlapping time windows. Panels B, C: whole brain correlates of resting-state BOLD with blink events and *GazeX* ^2^. Each analysis is corrected for multiple comparisons using FSL’s implementation of TFCE Family-wise-error control.

Whole brain correlations with eye-tracking metrics were found for the *blink f unction* and *GazeX* ^2^ regressors and presented in Figure 1B, C (*p <* .05, corrected for multiple comparisons with FWE; see Supplementary Table 1 for coordinates). We note these findings were identified via a Finite Impulse Response (FIR) analysis (see *Methods*) which estimated the HRF shape per regressor. Regressions based on canonical HRF-convolved regressors produced results that were not statistically significant.

### Eye tracking data: Correlation with Eye Orbit EPI data

We evaluated the correlation between each of the 12 types of eye tracking time series (see *Methods*) and the EO-EPI data. We controlled for the 12 tests using Bonferroni correction, because some of the tests were highly correlated (see Figure Appx.2). We found that three eye-tracking regressors significantly correlated with the EO-EPI envelope (Bonferroni corrected for 12 tests): the gaze power *vel GazeX* ^2^ + *vel GazeY* ^2^, square of pupil size *PupilSize*^2^, and the gaze velocity in the vertical (*Y*) direction. The pupil size was evaluated as deviation from the subject’s mean value, so its squared value indicated absolute deviations from mean value (we used squared deviation rather than absolute value as the derivative of the exponent is better behaved than that of the absolute function). Figure 2A shows sample time series reflecting raw EO-EPI, its envelope and eye-tracking regressors, and Figure 2B shows the estimated Kernels for gaze power and square of pupil size.

**Figure 2.**
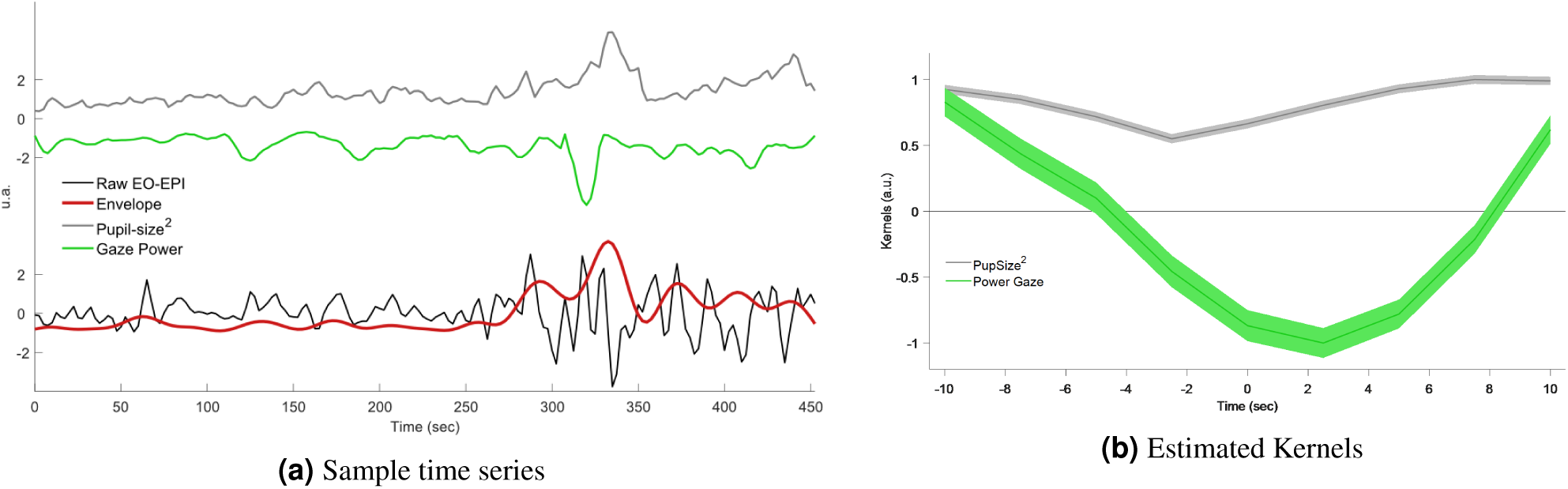
Relation between eye-tracking measures and EPI Orbit (EO-EPI) regressor.

Pupil-size squared explained 7 *±* 2% of the variance of the EO-EPI envelope and presented a significant positive correlation: *ρ* = 0.17 *±* 0.05, *t*(30) = 3.45, *p* = .0017, *d* = 0.62. Gaze power explained 5.4 *±* 1.6% of the variance of the EO-EPI envelope and had a significant negative correlation: *ρ* = −0.17 *±* 0.03, *t*(30) = 5.18, *p <* .001, *d* = 0.93. These two variables jointly explained the 11 *±* 3% of EO-EPI envelope’ variance, a significant improvement in model performance with respect the single variable cases (Δ*BIC <* −2). Gaze velocity in the *Y* direction had a weaker impact; it explained 3.7 *±* 1.0% of the EO-EPI’s envelope variance and had a significant positive correlation: *ρ* = 0.11 *±* 0.03, *t*(30) = 3.67, *p <* .001, *d* = 0.66. Adding this variable to the preceding regression model did not significantly increase explained variance (Δ*BIC* = −0.5). The exact numeric values corresponding to these kernels is given in Supplementary Table 2. Blinks were not significantly correlated with EO-EPI.

### Connectivity of EO-EPI regressors

We identified an extensive system that correlated with the EO-EPI regressor. For the convolved version of the EO-EPI regressor (EYE_*conv*_) we found correlations in pre- and post-central gyri bilaterally, parts of the superior temporal gyrus and visual cortex (Figure 3A). We also identified strong correlations (of opposite sign) in the thalamus (Figure 4A). We also found whole brain correlations for the non-convolved versions of the EO-EPI regressor (EYE_*raw*_). These were qualitatively similar, but reduced in extent (see Figures 3B, 4B). Whole-brain clusters for the EYE_*raw*_ and EYE_*conv*_ regressors are included in Supplementary Tables 3 and 4. A region of interest analysis indicated statistically significant correlations with EO-EPI in FEF (Wilcoxon *z* = 6.15, *p <* .001) but not in SEF (*z* = −1.28, *p >* .05).

**Figure 3.**
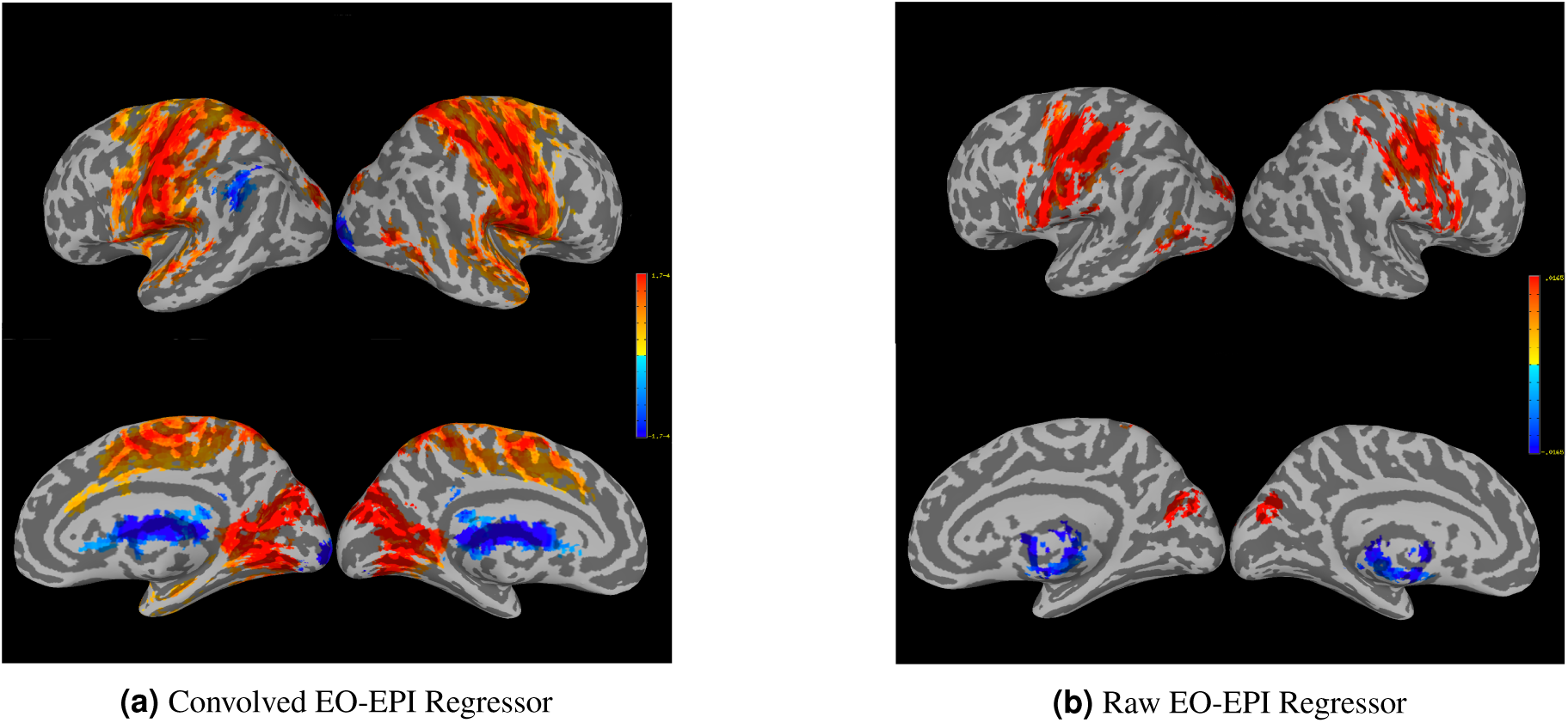
Whole-brain connectivity maps for the EYE_*conv*_ (Panel A) and EYE_*raw*_ regressors (Panel B).

**Figure 4.**
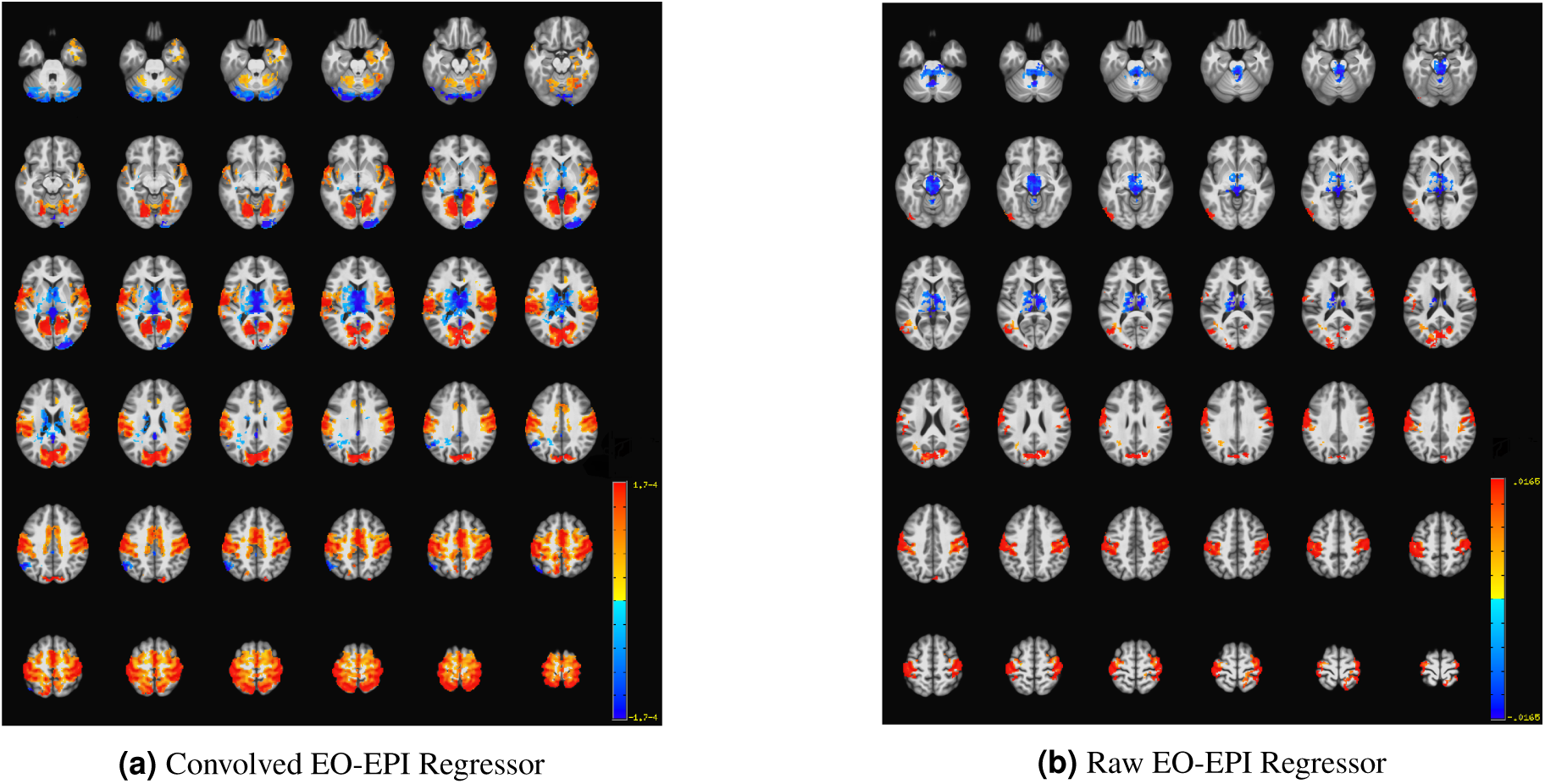
Axial slices showing whole-brain connectivity for the EYE_*conv*_ (Panel A) and EYE_*raw*_ regressors (Panel B).

In general, the tSNR of the raw time series was quite good across the cortex (see Figure Appx.5), with typical dropoff in low-signal and areas susceptible to motion. Values were similar to the those reported by the Human Connectome Project for 2mm and 3mm non-cleaned data (Smith et al., 2013). We treated each cluster where BOLD activity correlated with EO-EPI (raw or convolved) as a functional ROI and calculated the Mean and SD of tSNR in each cluster across participants. Most of these areas were associated with adequate tSNR, including the thalamus. This held for all statistically significant clusters picked up by the EYE_*raw*_ regressor (see Supplementary Table 5). For EYE_*conv*_ the clusters found in the left and right cerebellum were associated with low tSNR (and relatively systematically across participants, see Supplementary Table 6), as was a cluster in the mid occipital gyrus bilaterally (potentially as it includes time series from the field of view between the two hemisphere).

### EO-EPI regressor: between-participant differences in variance, power-spectra properties and relation to motion parameters

Across participants, the time series of the EO-EPI regressor presented a larger range of standard-deviation values than found in other ROIs. Figure 5A presents a histogram of the SD values for EYE_*raw*_ in the participant group, and comparative values from the temporoparietal junction (TPJ). The SD values for TP were relatively low and tightly clustered in the range of 5-45, with a mode of 10. In contrast, for the EO-EPI regressor, there was much less systematicity in the spread values across participants: the distribution of SD values was relatively more uniform and showed much larger values, some with *SD >* 200. The mean number of voxels in these regions was 1270 for TPJ and 406 for EYE_*raw*_.

**Figure 5.**
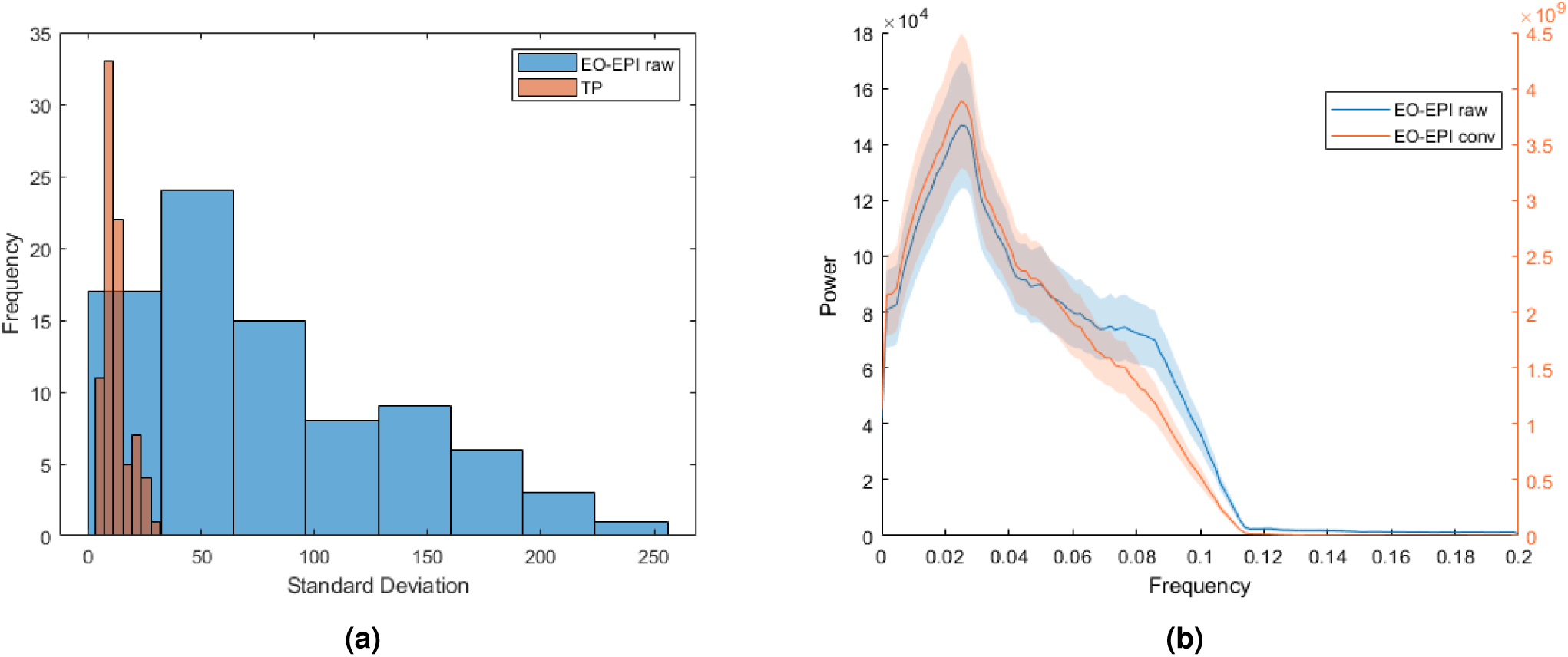
Spectral and spread-properties of EO-EPI regressor. Panel A: Across-participant distribution of standard deviations of EO-EPI time series and (for contrast) average time series from temporoparietal-junction ROI. Panel B: Frequency distribution of convolved and raw EO-EPI series. Differences in order of mangitude are due to convolution with HRF basis function.

The reason for these differences across participants is unclear. However, a byproduct is that when the EO-EPI regressor is correlated with brain activity in the context of regression, the resulting Beta values for this regressor have a very broad distribution with significant differences across participants and outliers. For this reason, using a parametric test on the group level can produce false-negatives or positives. To illustrate: in this current study, when non-parametric tests are used for group-level analysis, then both the Sign test and the Wilcoxon test produce group-level significance maps as reported here. AFNI’s multilevel analysis *3DMEMA* (G. Chen, Saad, Nath, Beauchamp, & Cox, 2012), which down-weights beta values from participants with noisier Beta estimates produces similar results, though statistically weaker. However, a typical group-level T-test of Beta values against zero produced a null result.

The large standard deviation of the EO-EPI regressor was related to peaks in that signal. As indicated in the *Methods* section, applying a ‘despiking’ procedure reduced the sensitivity of the whole brain correlation analysis: its most extreme effect was flattening several time series from the eye-orbit area, and in other cases it impacted a large number of time points in that area (see Figure Appx.3 for example). An analyses of the spectral features of EO-EPI (Figure 5B showed a strong peak in those time series at 0.04*Hz*, i.e., a cycle of 25sec. This is consistent with slow fluctuations sometimes seen in cortical regions. To summarize, the EO-EPI regressor, as would be expected, presented some time-domain features (spikes and strong inter-individual differences in spread) that differ from BOLD time series acquired in the brain and these need to be considered during pre-processing and group-level analyses. That said, its spectral power presented a strong peak at low frequencies of the sort seen in cortical BOLD time series.

With rare exceptions, EYE_*raw*_ was not-correlated with the estimated head-motion parameters. Significant correlations with any of the 6 motion parameters were found for 3 of the 83 participants: In the first case there was correlation with L/R displacement; in the second case there was correlation with L/R displacement and rotation; in the third case 5 of the 6 parameters were correlated. In all cases, correlation values were below 0.2. This lack of correlation suggests that variance in EYE_*raw*_ signal is not related to head motion, though an extreme case of movement may be picked up in this signal as well.

### Functional connectivity networks

An analysis of the network metrics revealed that several were significantly impacted by EO-EPI-removal, across all three Sparsity thresholds (i.e., top 10, 20 and 30% of connections). Difference values, effect sizes, and results of statistical tests are reported in Table 1. As shown in the Table, statistically significant results were associated with medium effect sizes in the range of 0.4-0.5. The raw connectivity matrices presented higher values for Maximal and Mean Node strength, Mean Cluster Coefficient (and transitivity). Conversely, maximized modularity was greater for the clean (EO-EPI-removed) matrices. Supplementary Table 7 reports the raw values for each metric.

**Table 1.** Difference of network metrics between Raw and Clean (EO-EPI-removed) functional connectivity matrices. Differences shown are in units of percentage apart from the number of communities and maximized modularity which maintain the original measure. *\*=p<*.*05, **=p<*.*005, ***=p<*.*001*

|  | Sparsity=0.1 |  |  | Sparsity=0.2 |  |  | Sparsity=0.3 |  |  |
| --- | --- | --- | --- | --- | --- | --- | --- | --- | --- |
|  | Difference | Cohen's D | T stat | Dif. | D | T | Dif | D | T |
| Max Degree | 1.23 | 0.38 | 3.40** | 0.74 | 0.37 | 3.33** | -0.04 | 0.02 | -0.22 |
| Min Degree | 3.45 | 0.04 | 0.38 | -2.58 | 0.08 | -0.76 | -1.90 | 0.10 | -0.92 |
| Max Strength | 2.45 | 0.49 | 4.41*** | 2.11 | 0.46 | 4.13*** | 1.91 | 0.42 | 3.79*** |
| Min Strength | 8.03 | 0.10 | 0.86 | -4.59 | 0.16 | -1.47 | -2.51 | 0.14 | -1.26 |
| Mean Strength | 0.99 | 0.49 | 4.36*** | 1.22 | 0.46 | 4.14*** | 1.38 | 0.44 | 3.94*** |
| Max Cluster Coefficient | 0.50 | 0.12 | 1.06 | 1.21 | 0.26 | 2.32* | 1.87 | 0.32 | 2.90** |
| Min Cluster Coefficient | 6.05 | 0.02 | 0.14 | -3.52 | 0.08 | -0.70 | -1.43 | 0.07 | -0.61 |
| Mean Cluster Coefficient | 1.08 | 0.46 | 4.11*** | 1.65 | 0.47 | 4.22*** | 1.84 | 0.44 | 3.94*** |
| Transitivity | 1.92 | 0.46 | 4.11*** | 2.39 | 0.46 | 4.14*** | 2.49 | 0.45 | 3.99*** |
| Assortativity | 0.31 | 0.06 | 0.54 | 1.44 | 0.22 | 1.95 | 2.12 | 0.28 | 2.46* |
| Efficiency | 0.12 | 0.18 | 1.60 | 0.58 | 0.49 | 4.39*** | 0.80 | 0.46 | 4.07*** |
| Max Number of Community | 0.02 | 0.01 | 0.10 | -0.02 | 0.05 | -0.42 | -0.02 | 0.05 | -0.44 |
| Maximized modularity | -0.007 | 0.44 | -3.95*** | -0.005 | 0.38 | -3.36** | -0.003 | 0.34 | -3.03** |
| Max betweenness centrality | -0.21 | 0.02 | -0.16 | 0.12 | 0.01 | 0.10 | 0.65 | 0.07 | 0.63 |
| Mean betweenness centrality | 0.86 | 0.41 | 3.70*** | 0.76 | 0.39 | 3.50*** | 0.25 | 0.14 | 1.29 |

Fitting the degree distributions using an exponentially truncated power law showed that the EO-EPI removed networks differed in the degree distribution (see Figure 6). As shown in the Figure, for 10% sparsity networks, EO-EPI removal impacted all three coefficients of the truncated power-law fit: coefficient: *t*(82) = 3.33, *p <* .01, *d* = 0.37, power law exponent, *t*(82) = −3.70, *p <* .001, *d* = 0.41, and degree cutoff point, *t*(82) = 3.59, *p <* .001, *d* = 0.4. For the 20% sparsity networks, differences were found for power law exponent, *t*(82) = −3.13, *p <* .01, *d* = 0.37, and degree cutoff point, *t*(82) = 2.59, *p <* .01, *d* = 0.33. No statistically significant differences were found for 30% sparsity networks. Figure Appx.6 presents mean degree-distributions for Raw and Clean networks in the different sparsity levels.

**Figure 6.**
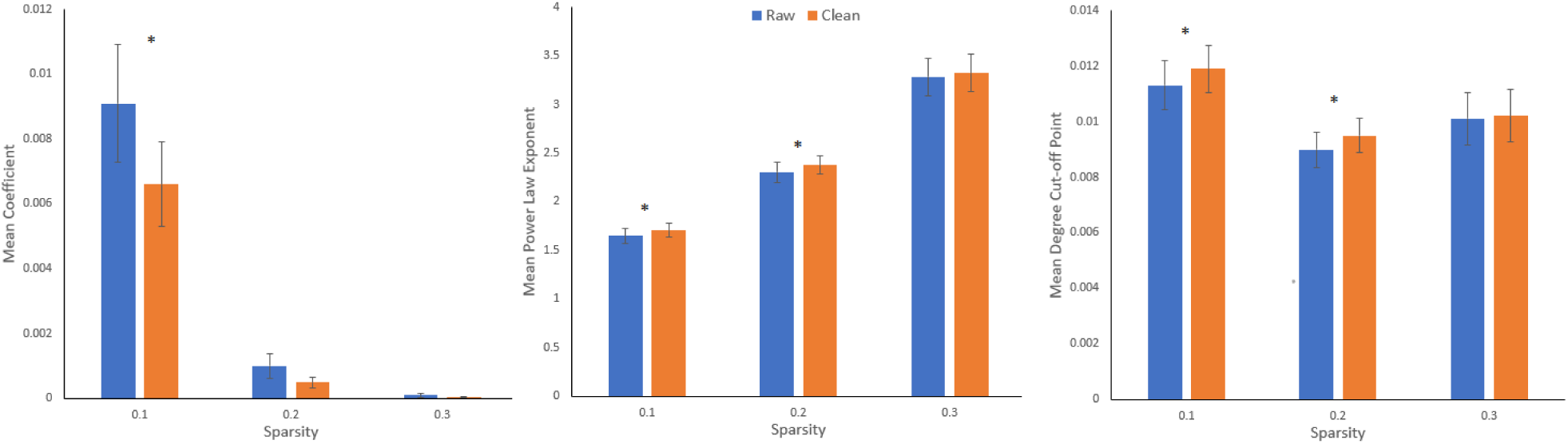
Mean power law parameters across sparsity levels (Coefficient, Power law exponent, Power law cutoff point). Bar-pairs for which a difference was significant are marked with a star (*).

We determined which areas tended to show changes in connectivity as function of EO-EPI removal. In general, this analysis is not independent of the whole-brain correlation with the EO-EPI time series used as a regressor, but it is more sensitive in identifying strongest pairwise differences. For each of the around 124,000 pairwise correlations we conducted a T-test to determine whether the pairwise correlations differed for raw and EO-EPI-removed connectivity matrices. The results (FDR corrected; Figure 7) showed that connectivity matrices constructed from the raw matrices presented stronger connectivity of sensory-motor areas with temporoparietal, dorsal-attention, visual cortex, and other sensory-motor regions. There were relatively few regions that showed stronger connectivity in the EO-EPI-removed condition, notably the Posterior Cingulate which showed stronger connectivity with multiple other brain areas.

**Figure 7.**
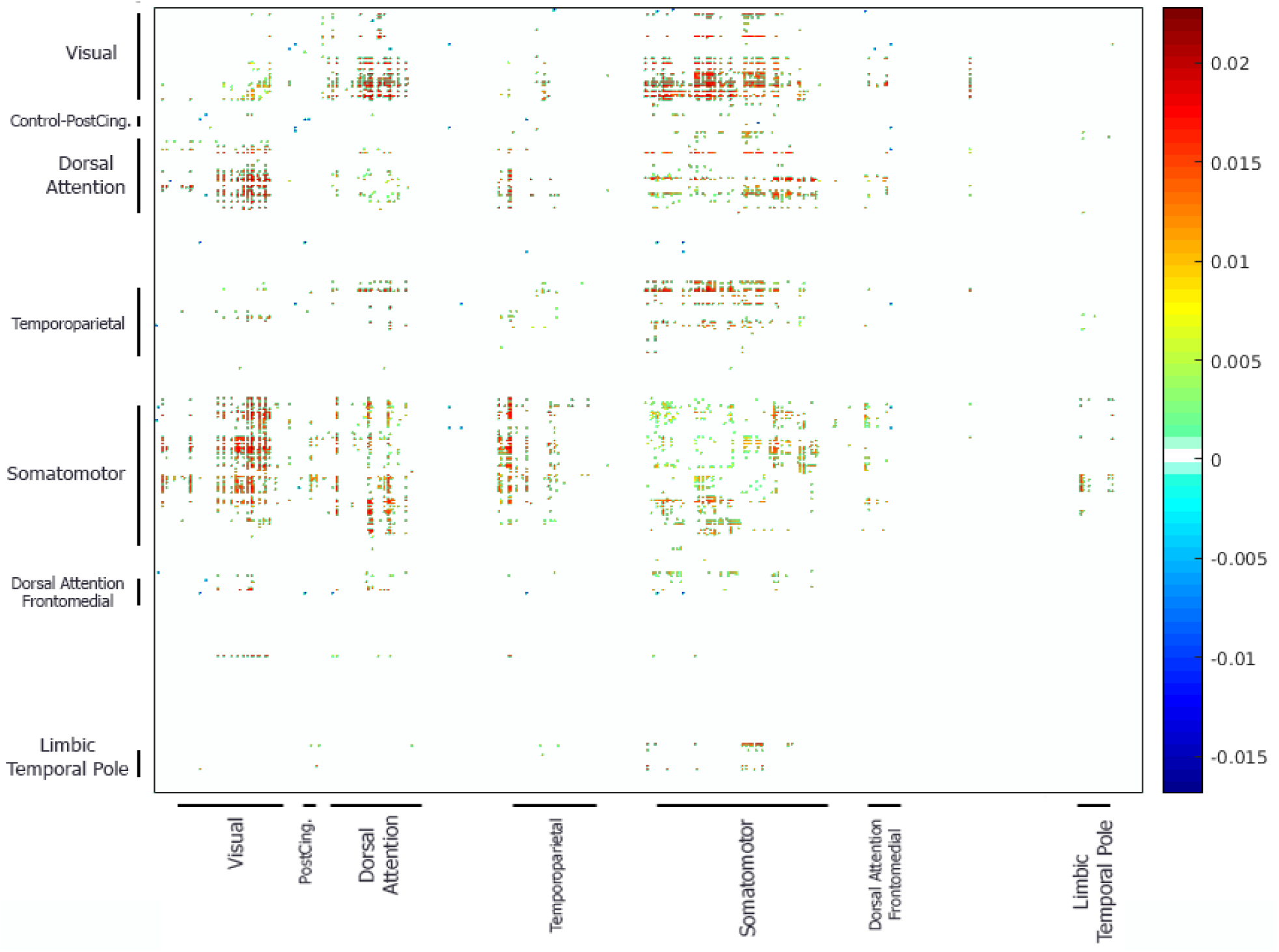
Pairwise-connectivity differences between raw and clean functional connectivity matrices (*Raw* − *clean, p <* 0.05, corrected for multiple comparisons with FDR).

The dual regression analysis did not identify any pre-defined RS network for which connectivity changed significantly. A hub-focused analysis that examined whether there were regions more frequently identified as hubs in the raw or EO-EPI-removed series also produced a null result: the most extreme example was a region defined as hub for 20 participants in one case and 25 in another (a non-significant difference on a binomial). While the location of these hubs was not a central point of the current study, broadly speaking, for the 10% sparsity threshold (raw) matrices, hubs were localized motor and sensory-motor areas (9 regions) Dorsal attention (6 regions), DMN (4 regions), temporal-parietal areas (4 regions) and ventral attention (2 areas). Only one visual extrastriate area was identified as a hub.

## Discussion

Neuroimaging is continuously expanding our understanding of the principles that determine structured patterns of RS connectivity. Our findings demonstrate that endogenous eye movements during RS contribute significantly to structured patterns of RS connectivity. Our main finding is that eye movements, measured via EPI time series recorded from the eye orbits, identified a sensory-motor system that appeared linked to oculomotor activity. Removal of activity accounted for by eye movements had systematic impact on whole-brain connectivity. We first address issues related to oculomotor measurement during the resting state that emerged in the study and then discuss the implications of the results for basic and applied research.

### Probing resting-state networks with Eye tracking and eye-orbit EPI data: technical considerations

As reviewed in the Introduction, few studies have studied brain activity patterns that are correlated with oculomotor activity during the resting state, and those have produced inconsistent and sometimes puzzling results. The most relevant is Fransson et al. (2014, N=18): It derived gaze-velocity data from eye tracking during a resting-state scan, finding correlation with DMN activity. Also related is McAvoy et al. (2012, N=9) which examined Brain/EOG correlations and reported a null result. In our own analyses of eye tracking data (N=32), we found correlation between BOLD-RS and only two eye tracking metrics: horizontal eye displacement, and blinks. These relatively modest correlations could be the result of noise in the eye tracking data, which presented itself in higher power across all frequencies for rejected data as compared to analyzed data.

We found correlations between the eye-tracking metrics and EPI data recorded from the eye orbit area (EO-EPI), Bonferroni corrected for 12 correlation tests. These were found for Gaze power, pupil size (squared), and gaze velocity in the Y direction. These data are consistent several prior reports. Beauchamp (2003) showed that peaks in the EO-EPI time series occur when an MR acquisition coincides with a rapid saccadic eye movement. Brodoehl, Witte, and Klingner (2016) and Son et al. (2019) showed that EO-EPI data can be used to estimate gaze location (when non-averaged; i.e., used in a multivariate context). In addition, Beauchamp’s observations suggest that for our interleaved acquisition, eye movements occurring either during odd- (up direction) or even-numbered (down direction) slice acquisition could be picked up in the analysis, because we treated the entire eye orbit as a single ROI. Consequently, while the volume acquisition time was 2.5sec, our effective temporal resolution for the eye-orbit ROI could have been higher, as we could identify eye-movement during both the up- or down-acquisition direction. EO-EPI fluctuations are likely mainly driven by signal disturbances due to air/tissue motion, but we cannot exclude the possibility that the signal also contains a BOLD component, due to the metabolic activity in nearby muscles. In particular, Law (1998) used PET rCBF to study brain systems involved in generation of voluntary saccades and reported active areas in the eye-orbit, “primary located close to the apex of the pyramidal shaped orbital cavity”. Our finding of a systematic delayed coupling in which changes in pupil size preceded EO-EPI fluctuations (the latter delayed by 2 sec), and of a strong peak frequency of 0.04Hz for EO-EPI are both consistent with the possibility that EO-EPI also reflects metabolic activity. We also found little independent evidence to suggest a strong contribution of motion artifacts to EO-EPI: beyond one participant for which 5 of 6 motion parameters correlated with EO-EPI, we only found 2 additional correlations with motion elements, for two additional participants.

Note that task compliance during this RS study was good. First, participants were continuously monitored and experimenters verified participants did not drift off to sleep during the scan. Second, the eye-tracking data indicated compliance with the task instructions in that the eye movements that were made during fixation were minor in magnitude (see Figure 1A). When evaluating average movements between successive 2sec epochs we found that in 75% of the cases, the magnitude was below 2degree, which corresponds to a small displacement. For this reason, we consider these data to be representative of typical compliant behavior during wakeful rest.

### Brain systems identified by Eye-Orbit EPI (EO-EPI) regressor

When used as a whole-brain regressor, the EO-EPI time series correlated with an extensive bilateral sensory-motor system. In addition, activity was found in superior parietal lobule, the dorsal part of the superior frontal gyrus, supplementary motor areas, and the extrastriate cortex in occipital lobe (excluding striate cortex). Region-of-interest analyses indicated activity in frontal eye fields. The topography of this system does not match either the ventral or dorsal attention networks as usually defined, but it is quite similar to the Frontal-Eye-Field connectivity map reported by Fox, Corbetta, Snyder, Vincent, and Raichle (2006). It is also highly similar to activity maps reported for simple eye movements in absence of attention, which have identified extensive activity in motor and premotor areas (e.g., Balslev et al., 2011) with little front-parietal involvement. A subset of these regions was also picked up by a non-convolved (‘Raw’) version of the EO-EPI regressor which may indicate that activity in these areas does not precede eye movements, but is relatively contemporaneous with them (to the extent that can be inferred from fMRI), or even that the eye movements reflected in the EO-EPI time series follow activity in those areas.

The brain areas we identify using EO-EPI (or eye tracking regressors) depart from ones frequently mentioned in studies of saccadic mechanisms, which prototypically reveal involvement of FEF/SEF and IPS. There are several possible explanations for this, which are not mutually exclusive. First, neuroimaging studies of saccades study saccade execution under exogenously determined conditions. Specifically, a distinction is made between two saccade categories, both externally-controlled: ‘reflexive’ saccades that orient to peripheral (typically sudden) target appearance, and ‘voluntary’ saccades that are not oriented towards a target in an unmediated manner but rather require a cognitive judgment prior to eye movement (for review, see Mort et al., 2003). These voluntary saccades are studied by paradigms such as anti-saccades (saccading to the opposite screen side of a target), memory-guided saccades (saccading to a location maintained in memory), or saccading to a location pre-cued by an arrow. Note that both reflexive and voluntary saccades are associated with few degrees of freedom with respect to the actual saccade-target, which constitutes a fundamental difference from the resting-state case. In addition, as indicated by Brown et al’ study (reviewed in the Introduction), activity in FEF/SEF/IPS may not be related to endogenous oculomotor control per se, but to the paradigm demands that require attention and detection of visual cues. In support of this possibility, a recent study (Agtzidis, MeyhÖ fer, Dorr, & Lencer, 2020) examining eye movements during naturalistic movie viewing similarly failed to identify a frontal parietal system related to saccades (neither dorsal nor ventral attention systems; see their Table 2), but instead documented saccade-related activity in visual cortex, and smooth-pursuit activity in precuneus, cingulate and occipital cortices. The authors attribute this failure to differences in paradigm, suggesting that natural viewing is associated with constant engagement rather than phasic shifts between fixation and saccades. This is also corroborated by the report by Son et al. (2019, N=5) showing that during naturalistic viewing, data acquired from the eye orbits correlates with brain activity in areas that do not resemble the topography of attentional networks (see their Figure 5).

Another possibility, which does not assume substantial differences between RS and active tasks, is technical in nature: it is possible that endogenous oculomotor-linked sensory motor activity during resting state is simply not often reported just because fixation is a frequently used implicit baseline in many oculomotor studies. If the network we identify is correlated with oculomotor activity both during fixation and saccade-to-target epochs (either reflexive or voluntary), then it will not be identifiable in analyses against baseline because it is partialled out as a result of that contrast.

### The impact of removal of EO-EPI properties from BOLD activity

We examined the impact of removing the variance related to EO-EPI from brain activity using a few well-defined topographical and topological properties. For topography we found that removal did not have a statistically significant impact on connectivity in any of the 14 well-defined resting state networks. We also examined the impact of removal on pair-wise regional connectivity using a 500-ROI parcellation (Schaefer et al., 2018). We grouped these 500 regions into 7 main clusters for purposes of graphical presentation (Figure 7). The analysis produced statistically significant effects (FDR corrected), mainly showing that EO-EPI-removal was associated with reduced connectivity between the somatomotor regions and visual, temporoparietal and also few dorsal-attention network areas. Also as shown in Figure 7, connectivity within each system was weakly impacted by EO-EPI removal if at all (i.e., few changes along the diagonal), which is consistent with the dual-regression results. To conclude, EO-EPI-removal appeared to primarily impact cross-network connectivity rather than within-network connectivity. Finally, we did not find evidence that EO-EPI-removal impacted the distribution of network-hubs in the brain.

However, robust results were found for both global and local topological metrics identified by a network analysis. We report only results that maintained across three sparsity thresholds: 10%, 20% and 30% of connections. For global properties, we find that modularity (Q) was higher for the clean matrices. For local properties, we found that the raw matrices were associated with greater node-strength values (indicating sum of connectivity linked to each node). For max-strength, the difference was 2.45% (effect size= 0.49). The mean cluster-coefficient (and strongly related, transitivity) were also impacted, showing reduced values (approaching 2.5% difference; effect-size=0.49) for the cleaned time series.

These changes are consistent with our other findings. EO-EPI is correlated with occipital, sensory-motor and few fronto-parietal areas, and as shown, EO-EPI removal predominantly impacts inter-regional / inter-internetwork connections rather than intra-network connections. For this reason, its removal serves to increase the modularity of resting state networks.

### Implications for network studies of typical and special populations

As indicated in a recent review (Hallquist & Hillary, 2018), graph theoretical approaches are increasingly applied in the context of resting-state fMRI studies of clinical disorders. In some cases, these features are deployed clinically to define new clinical subtypes, and in other cases, they are used to advance understanding of the brain systems that may be associated with the clinical deficit.

Being able to link differences in graph-theoretic-metrics to the oculomotor systems can increase the specificity of the explanations provided by RS analyses (by linking differences to a specific behavior), and would allow determining to what extent differences in RS connectivity between populations can be attributed to differences in oculomotor activity during resting-state acquisition.

A number of examples present the logic of this approach. For example, Parkinson’s Disease (PD) is associated with changes to functional connectivity when analyzed both from dynamic and static perspective (Kim et al., 2017). Neurophysiologically, it is associated with abnormality in eye movement control, including in generation of voluntary saccades. Anomalies are more evident for voluntary saccades, in early stages of disease (for review, see Pretegiani & Optican, 2017). A behavioral study (Zhang et al., 2018) showed that PD is linked to reduced fixation stability when fixation is required. Conversely, during free viewing of single images, PD patients make fewer saccadic eye movements, and within a more narrow range. Differences in network modularity for clinical populations have been documented in the case of Autism, which present lower modularity (Rudie et al., 2013) and Traumatic Brain Injury (Han et al., 2014) which has been associated with higher modularity and lower participation coefficient of sensory-motor systems (i.e., these areas are more weakly involved in between-module connectivity). In addition, both schizophrenia (e.g., Alexander-Bloch et al., 2012) and depression (e.g. Drysdale et al., 2017) have been linked to changes in RS connectivity. Alexander-Bloch et al. showed that schizophrenia is associated with reduced modularity in functional networks with motor areas bilaterally linked to different partitions, and Dysdale et al. used connectivity to identify four neurophysiological subtypes of depression based on functional connectivity, with two of the types showing markedly reduced connectivity (vs. ctrl) in nodes within sensory-motor systems we identified.

The findings could also have implications for the study of dynamic, time-varying connectivity in healthy and clinical populations. Knowing that some dynamic changes are associated with phasic states of eye movements would allow better interpretation of the drivers of time-varying dynamics. An early study of time-varying dynamics (Hutchison, Womelsdorf, Gati, Everling, & Menon, 2013) is consistent with this possibility. It documented time points presenting phasic, strong connectivity between frontal eye fields, sensory-motor regions and occipital regions, whereas such connectivity was completely absent at other time points. This suggests temporary synchronization of multiple brain networks in relation to eye movement.

### Conclusions

We found that oculomotor-movement provides a systematic contribution to RS connectivity in the human brain. It is correlated with activity in a brain network that largely involves sensory-motor and visual cortex, as well as the frontal eye fields. Removal of oculomotor contribution, as quantified via EPI time series sampled from the eye orbit area, produces changes to global topological features of RS networks. Isolating this contribution can produce a better understanding of activity sources that organize RS networks in health and disease, and could improve the use of RS network-features in the context of machine learning.

### Appendix

#### Supplementary Tables

**Table 1.**
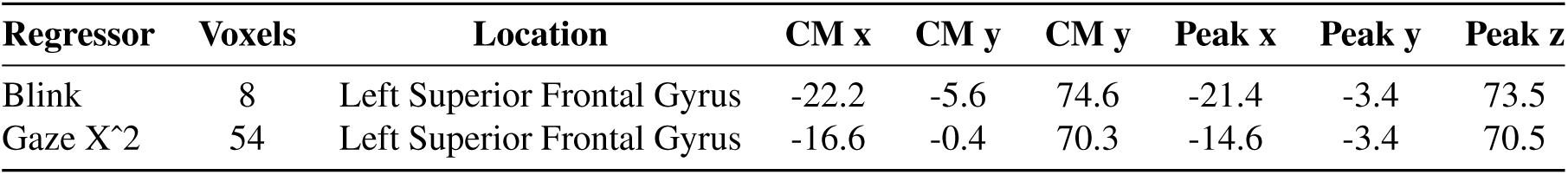
Cluster coordinates of the regions identified by the eye tracking data

| Regressor | Voxels | Location | CM x | CM y | CM y | Peak x | Peak y | Peak z |
| --- | --- | --- | --- | --- | --- | --- | --- | --- |
| Blink | 8 | Left Superior Frontal Gyrus | -22.2 | -5.6 | 74.6 | -21.4 | -3.4 | 73.5 |
| Gaze X^2 | 54 | Left Superior Frontal Gyrus | -16.6 | -0.4 | 70.3 | -14.6 | -3.4 | 70.5 |

**Table 2.** Numeric values describing kernels mediating the relationship between EO-EPI and eye tracking time series, for those eye tracking features for which the relation was statistically significant.

|  | Time in relation to peak value in eye-tracking series (sec) |  |  |  |  |  |  |  |  |
| --- | --- | --- | --- | --- | --- | --- | --- | --- | --- |
|  | -10 | -7.5 | -5 | -2.5 | 0 | 2.5 | 5 | 7.5 | 10 |
| Velocity of vertical gaze location | 0.146 | 0.245 | 0.313 | 0.338 | 0.328 | 0.280 | 0.205 | 0.117 | 0.031 |
| Pupil size squared | 0.101 | 0.093 | 0.078 | 0.060 | 0.073 | 0.088 | 0.101 | 0.109 | 0.108 |
| Velocity of Gaze Amplitude | 0.027 | 0.014 | 0.003 | -0.015 | -0.029 | -0.033 | -0.026 | -0.007 | 0.020 |

**Table 3.** Cluster mass and peak coordinates of the regions by *EYE*_*conv*_.

| Voxels | Location | CM x | CM y | CM z | Peak x | Peak y | Peakz | Mean Delta | Mean Z |
| --- | --- | --- | --- | --- | --- | --- | --- | --- | --- |
| 18517 | Right Cingulate Gyrus | -4.1 | 21.4 | 40.0 | -37.1 | 25.9 | 70.5 | 0.00012 | 3.41 |
| 5144 | Right Lingual Gyrus | -2.0 | 68.6 | 7.5 | 10.1 | 66.4 | -1.5 | 0.000151 | 3.23 |
| 2521 | Left Thalamus | 2.1 | 19.4 | 12.4 | 1.1 | 23.6 | 13.5 | -0.000117 | -3.29 |
| 1806 | Left Cerebellum/Declive | 4.3 | 79.7 | -29.5 | -16.9 | 84.4 | -19.5 | -0.00011 | -3.039 |
| 668 | Right Cerebellum/Inferior Semi-Lunar Lobule | -21.1 | 62.5 | -52.5 | -28.1 | 52.9 | -52.5 | 0.00007 | 3.31 |
| 592 | Left Inferior Parietal Lobule | 45.1 | 55.7 | 38.3 | 57.4 | 59.6 | 43.5 | -0.000094 | -2.92 |
| 547 | Middle Occipital Gyrus | -21.8 | 96.3 | -0.9 | -12.4 | 1.5 | 100.1 | -0.00015 | -3.15 |
| 488 | Left Cerebellum/Inferior Semi-Lunar Lobule | 21.7 | 62.8 | -52.9 | 21.4 | 64.1 | -52.5 | 0.000068 | 3.21 |

**Table 4.** Cluster mass and peak coordinates of the regions that are identified by *EYE*_*raw*_.

| Voxels | Location | CM x | CM y | CM y | Peak x | Peak y | Peak z | Mean Delta | Mean Z |
| --- | --- | --- | --- | --- | --- | --- | --- | --- | --- |
| 2888 | Cerebellar Vermis | 0.1 | 27.9 | -12.8 | 3.4 | 43.9 | 4.5 | -0.01313 | -3.12 |
| 1937 | Right Postcentral Gyrus | -45.8 | 19.2 | 49.7 | -39.4 | 21.4 | 70.5 | 0.01696 | 3.04 |
| 1901 | Left Postcentral Gyrus | 50.3 | 20.7 | 47 | 43.9 | 21.4 | 67.5 | 0.018062 | 3.07 |
| 631 | Left Cuneus | 0.6 | 79.9 | 23.4 | -1.1 | 79.9 | 37.5 | 0.019491 | 2.90 |
| 517 | Left Middle Occipital Gyrus | 47.1 | 70.5 | 3.8 | 52.9 | 70.9 | 13.5 | 0.014119 | 2.80 |

**Table 5.** Mean tSNR values of the clusters identified by *EYE*_*raw*_

| Location | Mean tSNR | SD of tSNR |
| --- | --- | --- |
| Cerebellar Vermis | 62.58 | 21.35 |
| Right Postcentral Gyrus | 68.74 | 29.89 |
| Left Postcentral Gyrus | 66.8 | 29.43 |
| Left Cuneus | 78.19 | 34.38 |
| Left Middle Occipital Gyrus | 78.01 | 32.49 |

**Table 6.** Mean tSNR values of the clusters identified by *EYE*_*conv*_.

| Location | Mean tSNR | SD of tSNR |
| --- | --- | --- |
| Right Cingulate Gyrus | 67.75 | 28.96 |
| Right Lingual Gyrus | 70.06 | 30.53 |
| Left Thalamus | 61.26 | 22.71 |
| Left Cerebellum/Declive | 45.87 | 25.49 |
| Right Cerebellum/Inferior Semi-Lunar Lobule | 48.76 | 25.30 |
| Left Inferior Parietal Lobule | 86.02 | 32.38 |
| Middle Occipital Gyrus | 45.10 | 25.21 |
| Left Cerebellum/Inferior Semi-Lunar Lobule | 44.55 | 25.29 |

**Table 7.** Means for network metrics for Raw and Clean (EO-EPI-removed) functional connectivity matrices in different sparsity levels.

|  | Sparsity=0.1 |  | Sparsity=0.2 |  | Sparsity=0.3 |  |
| --- | --- | --- | --- | --- | --- | --- |
|  | Mean raw | Mean clean | Mean raw | Mean clean | Mean raw | Mean clean |
| Max Degree | 146.03 | 144.23 | 220.88 | 219.23 | 278.88 | 279.02 |
| Min Degree | 0.36 | 0.35 | 3.863 | 3.963 | 13.113 | 13.363 |
| Max Strength | 91.07 | 88.84 | 119.66 | 117.13 | 135.16 | 132.57 |
| Min Strength | 0.18 | 0.17 | 1.457 | 1.524 | 3.934 | 4.033 |
| Mean Strength | 29.74 | 29.45 | 50.09 | 49.48 | 65.367 | 64.464 |
| Max Cluster Coefficient | 0.672 | 0.668 | 0.529 | 0.522 | 0.442 | 0.434 |
| Min Cluster Coefficient | 0.004 | 0.003 | 0.061 | 0.063 | 0.111 | 0.112 |
| Mean Cluster Coefficient | 0.339 | 0.336 | 0.308 | 0.303 | 0.283 | 0.278 |
| Transitivity | 0.359 | 0.352 | 0.323 | 0.315 | 0.296 | 0.289 |
| Assortativity | 0.4220 | 0.4207 | 0.354 | 0.349 | 0.305 | 0.298 |
| Efficiency | 0.262 | 0.261 | 0.287 | 0.285 | 0.292 | 0.289 |
| Max Number of Community | 5.57 | 5.55 | 3.938 | 3.963 | 3.475 | 3.500 |
| Maximized Modularity | 0.44 | 0.45 | 0.353 | 0.358 | 0.300 | 0.303 |
| Max Betweenness Centrality | 6979.50 | 6994.27 | 4529.85 | 4524.27 | 4086.70 | 4059.82 |
| Min Betweenness Centrality | 773.09 | 766.48 | 537.955 | 533.852 | 475.344 | 474.124 |

#### Supplementary Figures

**Figure Appx.1.**
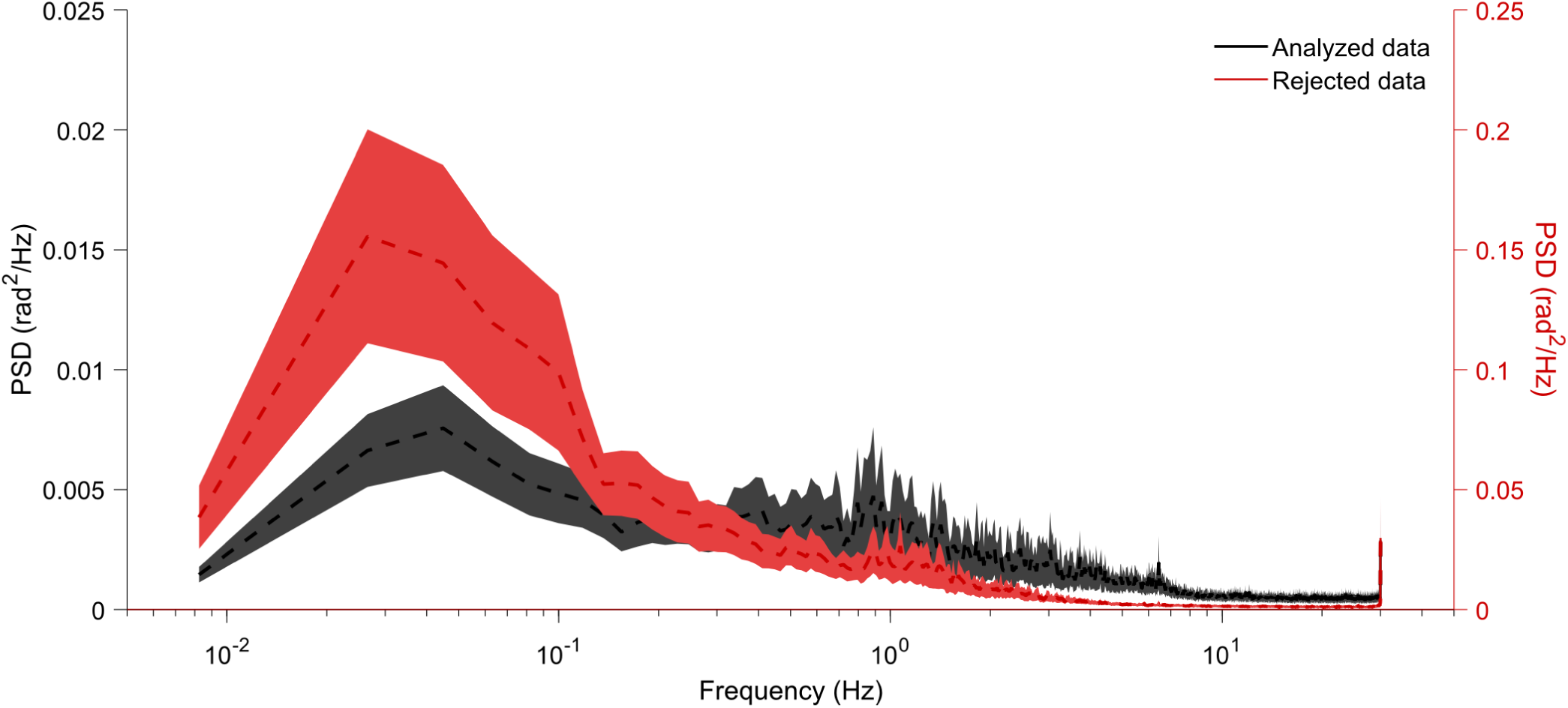
Power spectra of eye-tracking data for data rejected or maintained by the quality-control procedure. Note the dual Y-axis scales. Rejected data presented power around one order of magnitude higher than maintained data. This difference was significant even for the highest measurable frequency, where power in rejected data was 2.5 times higher than that maintained.

**Figure Appx.2.**
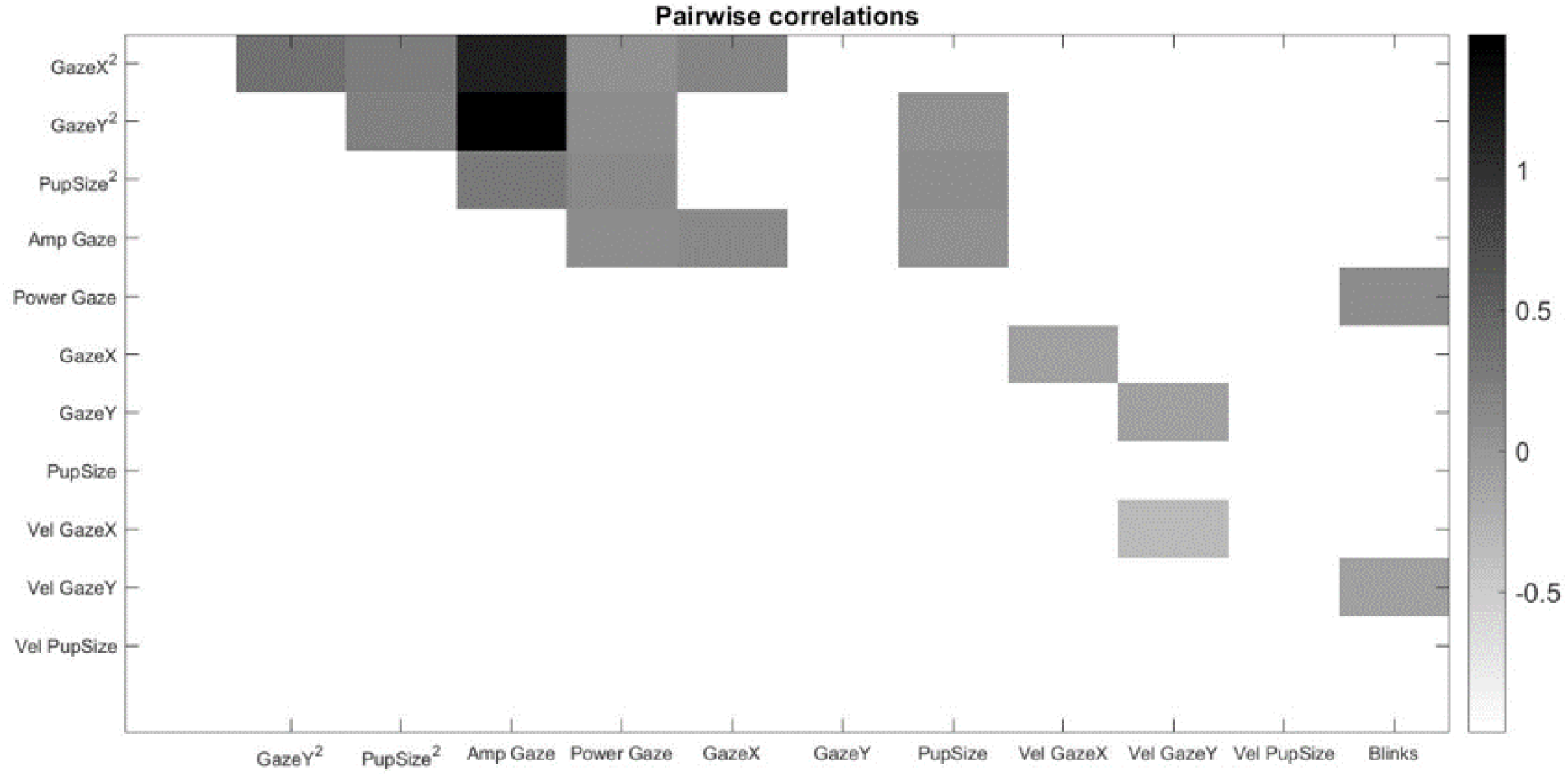
Correlation between different eye-tracking metrics. Only statistically significant correlations shown. Correlation between Gaze Power and *Gaze*^2^ or *Gaze*^2^ reflects a spurious correlation as Gaze Power is the sum of those two terms.

**Figure Appx.3.**
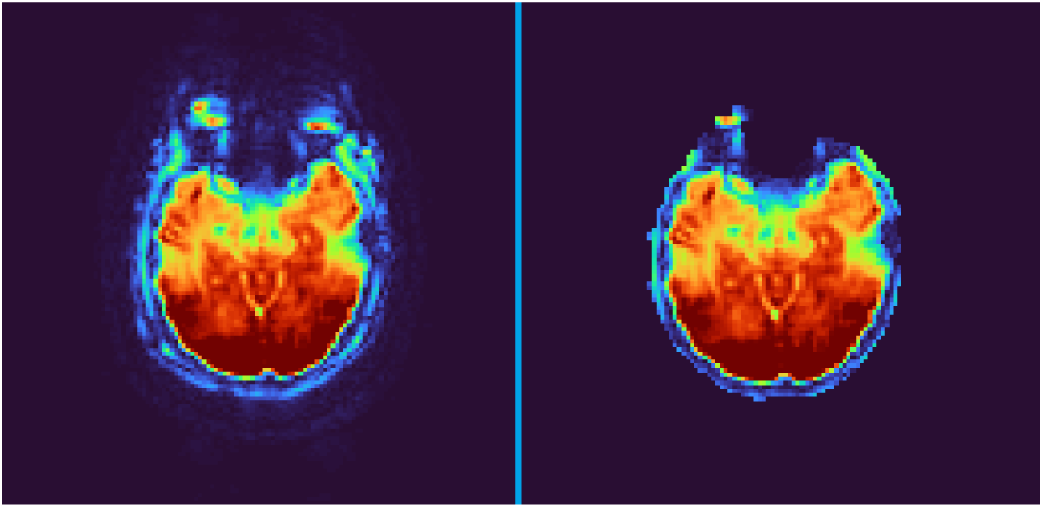
Impact of despiking on remaining signal in Eye Orbit area. Left image: without despiking. Right image: despiked.

**Figure Appx.4.**
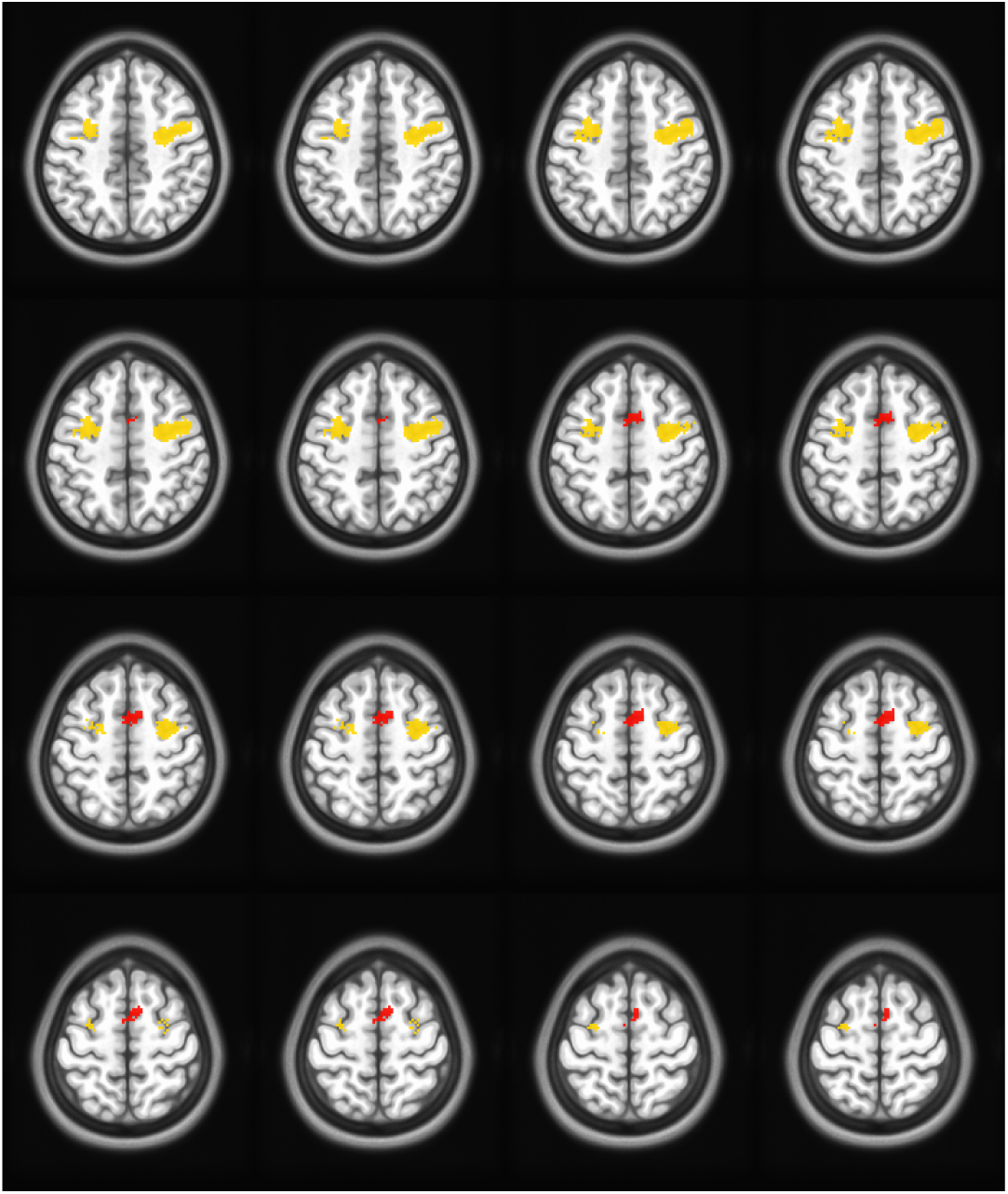
FEF (yellow) and SEF (red) ROIs derived from Neurosynth.

**Figure Appx.5.**
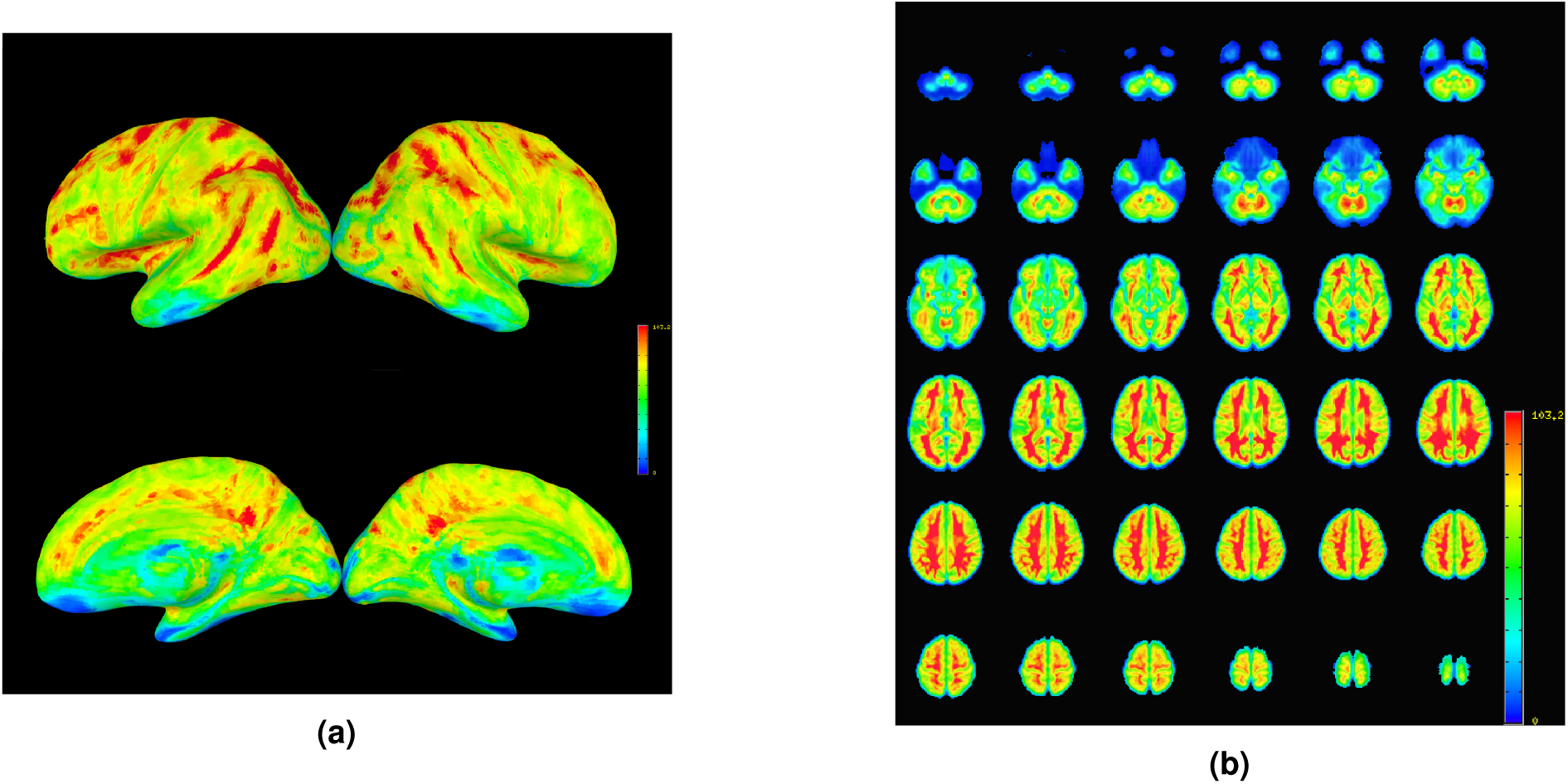
Temporal Signal-To-Noise (tSNR) values presented on Axial (a) and Surface (b) representations. These were calculated as *Mean/SD* of the raw time series prior to any pre-processing

**Figure Appx.6.**
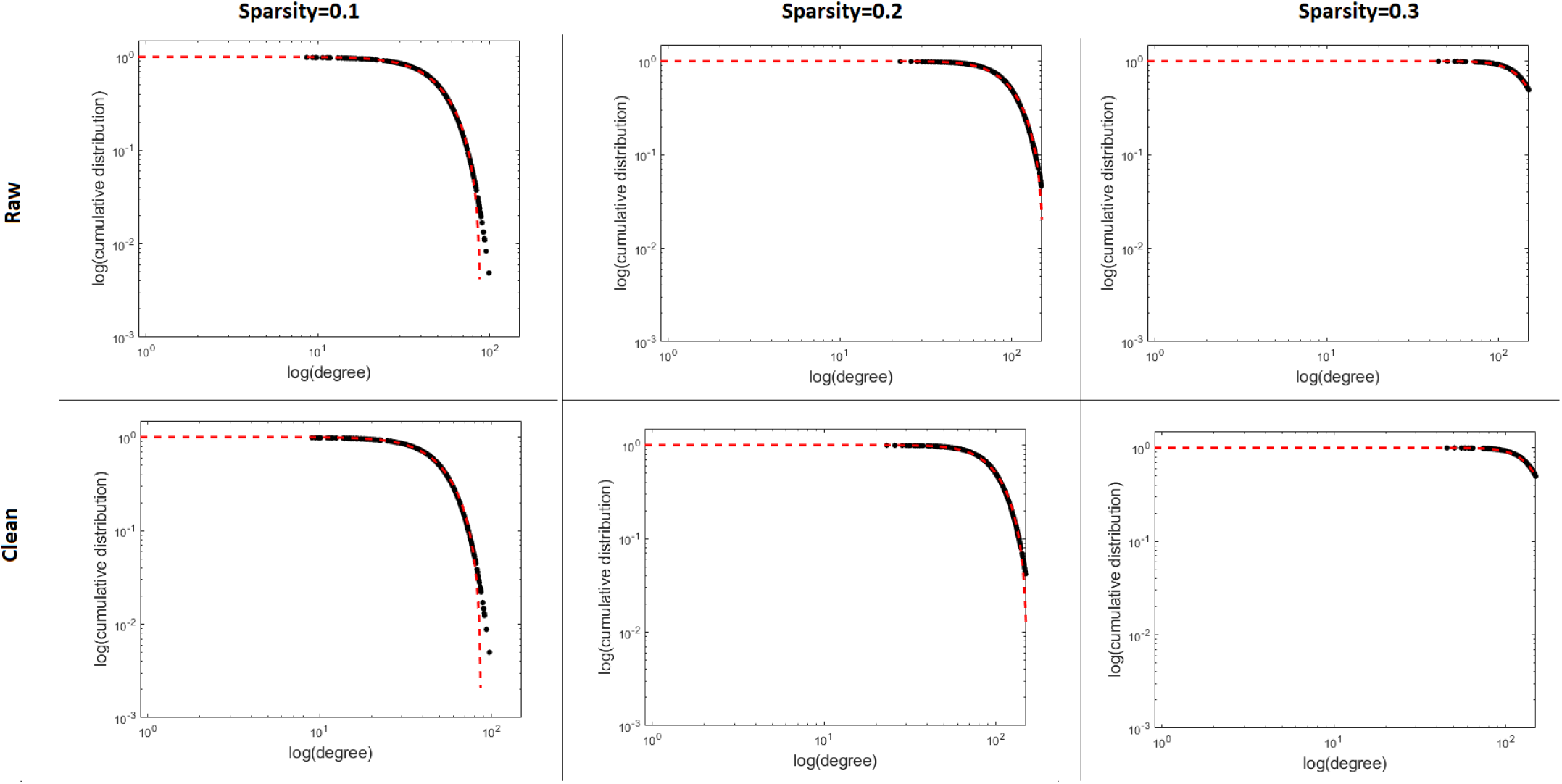
Mean degree distribution of raw adn clean matrices across sparsity levels.

#### Description of chosen network metrics

- **Degree**: Node degree is the number of links connected to the node. In directed networks, the in-degree is the number of inward links and the out-degree is the number of outward links. Connection weights are ignored in calculations.
- **Strength**: Node strength is the sum of weights of links connected to the node. In directed networks, the in-strength is the sum of inward link weights and the out-strength is the sum of outward link weights.
- **Clustering coefficient**: The clustering coefficient is the fraction of triangles around a node and is equivalent to the fraction of node’s neighbors that are neighbors of each other.
- **Transitivity**: The transitivity is the ratio of triangles to triplets in the network and is an alternative to the clustering coefficient.
- **Assortativity**: The assortativity coefficient is a correlation coefficient between the degrees of all nodes on two opposite ends of a link. A positive assortativity coefficient indicates that nodes tend to link to other nodes with the same or similar degree.
- **Global Efficiency**: The global efficiency is the average inverse shortest path length in the network, and is inversely related to the characteristic path length.
- **Number of communities**: The optimal community structure is a subdivision of the network into nonoverlapping groups of nodes in a way that maximizes the number of within-group edges, and minimizes the number of between-group edges. To calculate maximum number of community for each network, we recorded the largest number of community.
- **Modularity**: The modularity is a statistic that quantifies the degree to which the network may be subdivided into such clearly delineated groups.
- **Betweenness Centrality**: Node betweenness centrality is the fraction of all shortest paths in the network that contains a given node. Nodes with high values of betweenness centrality participate in a large number of shortest paths

## Acknowledgments

The authors are grateful to the Sleepy Brain project team for openly sharing the data, to staff physicist Rouslan Sitnikov for technical advice, and to Jorge Jovicich for assistance in assessment of ghosting artifacts.

## Notes

1 This mis-measurement is known as the Pupil Foreshortening Error (PFE). For example, Hayes and Petrov (2016) showed that deviations from center of camera-view produce systematic PFEs that can reach *∼*12% at typical viewing distances. Significant PFEs were produced even with movements as small as *∼*4*°* of center.

2 All data available online at https://www.openneuro.org/datasets/ds000201/

3 *Pupil size* was defined as (*pupil width* + *pupil height*)*/*2; we note that with our instrumentation (as well as many other eye trackers), pupil size is confounded with gaze position (Hayes & Petrov, 2016), resulting in significant correlations between *pupil size* and the gaze both in *x* and *y* directions (*p <* .01 in 30/32 subjects).

## References

Agtzidis, I., Meyhöfer, I., Dorr, M., & Lencer, R. (2020). Following forrest gump: Smooth pursuit related brain activation during free movie viewing. NeuroImage, 116491.

Alexander-Bloch, A., Lambiotte, R., Roberts, B., Giedd, J., Gogtay, N., & Bullmore, E. (2012). The discovery of population differences in network community structure: New methods and applications to brain functional networks in schizophrenia. Neuroimage, 59(4), 3889–3900.

Avants, B. B., Tustison, N., & Song, G. (2011). Advanced Normalization Tools (ANTS) Brian B. Avants, Nick Tustison and Gang Song, 1–35. Retrieved from www.picsl.upenn.edu/ANTS.

Balslev, D., Albert, N. B., & Miall, C. (2011). Eye muscle proprioception is represented bilaterally in the sensorimotor cortex. Human brain mapping, 32(4), 624–631.

Beauchamp, M. S. (2003). Detection of eye movements from fmri data. Magnetic Resonance in Medicine, 49(2), 376–380. Read. doi:10.1002/mrm.10345

Becker, W., & Fuchs, A. (1969). Further properties of the human saccadic system: Eye movements and correction saccades with and without visual fixation points. Vision Research, 9(10), 1247–1258. read. doi:10.1016/0042-6989(69)90112-6

Beckmann, C., Mackay, C., Filippini, N., & Smith, S. (2009). Group comparison of resting-state FMRI data using multi-subject ICA and dual regression. NeuroImage. doi:10.1016/s1053-8119(09)71511-3

Birn, R. M. (2012). The role of physiological noise in resting-state functional connectivity. Neuroimage, 62(2), 864–870.

Brodoehl, S., Witte, O. W., & Klingner, C. M. (2016). Measuring eye states in functional mri. BMC Neuroscience, 17(1). Read. doi:10.1186/s12868-016-0282-7

Brown, M. R., Goltz, H. C., Vilis, T., Ford, K. A., & Everling, S. (2006). Inhibition and generation of saccades: Rapid event-related fmri of prosaccades, antisaccades, and nogo trials. NeuroImage, 33(2), 644–659. doi:10.1016/j.neuroimage.2006.07.002

Chen, G., Saad, Z. S., Nath, A. R., Beauchamp, M. S., & Cox, R. W. (2012). Fmri group analysis combining effect estimates and their variances. Neuroimage, 60(1), 747–765.

Chen, J., Lewis, L., Chang, C., Tian, Q., Fultz, N., Ohringer, N., … Polimeni, J. (2020). Resting-state “physiological networks”. NeuroImage, 116707.

Chen, W., & Zhu, X.-H. (1997). Suppression of physiological eye movement artifacts in functional mri using slab presaturation. Magnetic Resonance in Medicine, 38(4), 546–550. Read. doi:10.1002/mrm.1910380407

Cox, R. W. (1996). AFNI: Software for analysis and visualization of functional magnetic resonance neuroimages. Computers and Biomedical Research, 29(3), 162–173. doi:10.1006/cbmr.1996.0014

de Vos, F., Koini, M., Schouten, T. M., Seiler, S., van der Grond, J., Lechner, A., … Rombouts, S. A. (2018). A comprehensive analysis of resting state fmri measures to classify individual patients with alzheimer’s disease. Neuroimage, 167, 62–72.

Drysdale, A. T., Grosenick, L., Downar, J., Dunlop, K., Mansouri, F., Meng, Y., … Etkin, A., et al. (2017). Resting-state connectivity biomarkers define neurophysiological subtypes of depression. Nature medicine, 23(1), 28.

Dubois, J., Galdi, P., Paul, L. K., & Adolphs, R. (2018). A distributed brain network predicts general intelligence from resting-state human neuroimaging data. Philosophical Transactions of the Royal Society B: Biological Sciences, 373(1756), 20170284.

Fonov, V., Evans, A., McKinstry, R., Almli, C., & Collins, D. (2009). Unbiased nonlinear average age-appropriate brain templates from birth to adulthood. NeuroImage. doi:10.1016/s1053-8119(09)70884-5

Fornito, A., Zalesky, A., & Bullmore, E. T. (2010). Network scaling effects in graph analytic studies of human resting-state fmri data. Frontiers in systems neuroscience, 4, 22.

Fox, M. D., Corbetta, M., Snyder, A. Z., Vincent, J. L., & Raichle, M. E. (2006). Spontaneous neuronal activity distinguishes human dorsal and ventral attention systems. Proceedings of the National Academy of Sciences, 103(26), 10046–10051.

Fransson, P., Flodin, P., Seimyr, G. Ö., & Pansell, T. (2014). Slow fluctuations in eye position and resting-state functional magnetic resonance imaging brain activity during visual fixation. European Journal of Neuroscience, 40(12), 3828–3835. Read. doi:10.1111/ejn.12745

Hallquist, M. N., & Hillary, F. G. (2018). Graph theory approaches to functional network organization in brain disorders: A critique for a brave new small-world. Network Neuroscience, 3(1), 1–26.

Han, K., Mac Donald, C. L., Johnson, A. M., Barnes, Y., Wierzechowski, L., Zonies, D., … Raichle, M. E., et al. (2014). Disrupted modular organization of resting-state cortical functional connectivity in us military personnel following concussive ‘mild’blast-related traumatic brain injury. Neuroimage, 84, 76–96.

Hayes, T. R., & Petrov, A. A. (2016). Mapping and correcting the influence of gaze position on pupil size measurements. Behavior Research Methods, 48(2), 510–527. read. doi:10.3758/s13428-015-0588-x

Honey, C., Sporns, O., Cammoun, L., Gigandet, X., Thiran, J.-P., Meuli, R., & Hagmann, P. (2009). Predicting human resting-state functional connectivity from structural connectivity. Proceedings of the National Academy of Sciences, 106(6), 2035–2040.

Hutchison, R. M., Womelsdorf, T., Gati, J. S., Everling, S., & Menon, R. S. (2013). Resting-state networks show dynamic functional connectivity in awake humans and anesthetized macaques. Human brain mapping, 34(9), 2154–2177.

Jenkinson, M., Beckmann, C. F., Behrens, T. E., Woolrich, M. W., & Smith, S. M. (2012). Fsl. Neuroimage, 62(2), 782–790. doi:10.1016/j.neuroimage.2011.09.015. arXiv:1401.4122v2

Kelly, A. C., Uddin, L. Q., Biswal, B. B., Castellanos, F. X., & Milham, M. P. (2008). Competition between functional brain networks mediates behavioral variability. Neuroimage, 39(1), 527–537.

Kim, J., Criaud, M., Cho, S. S., Diez-Cirarda, M., Mihaescu, A., Coakeley, S., … Houle, S., et al. (2017). Abnormal intrinsic brain functional network dynamics in parkinson’s disease. Brain, 140(11), 2955–2967.

Law, I. (1998). Parieto-occipital cortex activation during self-generated eye movements in the dark. Brain, 121(11), 2189–2200. read1. doi:10.1093/brain/121.11.2189

Marx, E., Stephan, T., Nolte, A., Deutschländer, A., Seelos, K. C., Dieterich, M., & Brandt, T. (2003). Eye closure in darkness animates sensory systems. Neuroimage, 19(3), 924–934.

McAvoy, M., Larson-Prior, L., Ludwikow, M., Zhang, D., Snyder, A. Z., Gusnard, D. L., … d’Avossa, G. (2012). Dissociated mean and functional connectivity bold signals in visual cortex during eyes closed and fixation. Journal of neurophysiology, 108(9), 2363–2372.

Mišić, B., Betzel, R. F., De Reus, M. A., Van Den Heuvel, M. P., Berman, M. G., McIntosh, A. R., & Sporns, O. (2016). Network-level structure-function relationships in human neocortex. Cerebral Cortex, 26(7), 3285–3296.

Mort, D. J., Perry, R. J., Mannan, S. K., Hodgson, T. L., Anderson, E., Quest, R., … Kennard, C. (2003). Differential cortical activation during voluntary and reflexive saccades in man. NeuroImage, 18(2), 231–246. read. ok intro. doi:10.1016/S1053-8119(02)00028-9

Nickerson, L. D., Smith, S. M., Öngür, D., & Beckmann, C. F. (2017). Using dual regression to investigate network shape and amplitude in functional connectivity analyses. Frontiers in Neuroscience. doi:10.3389/fnins.2017.00115

Nilsonne, G., Tamm, S., D’Onofrio, P., Thuné, H. Å., Schwarz, J., Lavebratt, C., … Åkerstedt, T. (2016). A multimodal brain imaging dataset on sleep deprivation in young and old humans, 1–27. Retrieved from https://openarchive.ki.se/xmlui/handle/10616/45181

Nostro, A. D., Müller, V. I., Varikuti, D. P., Pläschke, R. N., Hoffstaedter, F., Langner, R., … Eickhoff, S. B. (2018). Predicting personality from network-based resting-state functional connectivity. Brain Structure and Function, 223(6), 2699–2719.

Parkes, L., Fulcher, B., Yücel, M., & Fornito, A. (2018). An evaluation of the efficacy, reliability, and sensitivity of motion correction strategies for resting-state functional mri. Neuroimage, 171, 415–436.

Pretegiani, E., & Optican, L. M. (2017). Eye movements in parkinson’s disease and inherited parkinsonian syndromes. Frontiers in neurology, 8, 592.

Ramot, M., Wilf, M., Goldberg, H., Weiss, T., Deouell, L. Y., & Malach, R. (2011). Coupling between spontaneous (resting state) fMRI fluctuations and human oculo-motor activity. NeuroImage, 58(1), 213–225. doi:10.1016/j.neuroimage.2011.06.015

Rorden, C., Karnath, H. O., & Bonilha, L. (2007). Improving lesion-symptom mapping. Journal of Cognitive Neuroscience. doi:10.1162/jocn.2007.19.7.1081

Rosenberg, M. D., Hsu, W.-T., Scheinost, D., Constable, T. R., & Chun, M. M. (2018). Connectome-based models predict separable components of attention in novel individuals. Journal of Cognitive Neuroscience, 30(2), 160–173.

Rubinov, M., & Sporns, O. (2010). Complex network measures of brain connectivity: Uses and interpretations. NeuroImage. doi:10.1016/j.neuroimage.2009.10.003

Rudie, J. D., Brown, J., Beck-Pancer, D., Hernandez, L., Dennis, E., Thompson, P., … Dapretto, M. (2013). Altered functional and structural brain network organization in autism. NeuroImage: clinical, 2, 79–94.

Schaefer, A., Kong, R., Gordon, E. M., Laumann, T. O., Zuo, X.-N., Holmes, A. J., … Yeo, B. T. T. (2018). Local-Global Parcellation of the Human Cerebral Cortex from Intrinsic Functional Connectivity MRI. Cerebral Cortex. doi:10.1093/cercor/bhx179

Shirer, W. R., Ryali, S., Rykhlevskaia, E., Menon, V., & Greicius, M. D. (2012). Decoding subject-driven cognitive states with whole-brain connectivity patterns. Cerebral Cortex. doi:10.1093/cercor/bhr099

Siegel, S., & Castellan, N. J. (1956). Nonparametric statistics for the behavioral sciences. McGraw-hill New York.

Smith, S. M., Beckmann, C. F. [Christian F], Andersson, J., Auerbach, E. J., Bijsterbosch, J., Douaud, G., … Harms, M. P., et al. (2013). Resting-state fmri in the human connectome project. Neuroimage, 80, 144–168.

Son, J., Ai, L., Lim, R., Xu, T., Colcombe, S., Franco, A. R., … Milham, M. (2019). Evaluating fmri-based estimation of eye gaze during naturalistic viewing. Cerebral Cortex. doi:10.1093/cercor/bhz157

Takarae, Y., Minshew, N. J., Luna, B., Krisky, C. M., & Sweeney, J. A. (2004). Pursuit eye movement deficits in autism. Brain, 127(12), 2584–2594.

West, G. L., Welsh, T. N., & Pratt, J. (2009). Saccadic trajectories receive online correction: Evidence for a feedback-based system of oculomotor control. Journal of Motor Behavior, 41(2), 117–127.

Xu, P., Huang, R., Wang, J., Van Dam, N. T., Xie, T., Dong, Z., … jia Luo, Y. (2014). Different topological organization of human brain functional networks with eyes open versus eyes closed. NeuroImage, 90, 246–255. doi:10.1016/j.neuroimage.2013.12.060

Yarkoni, T., Poldrack, R. A., Nichols, T. E., Van Essen, D. C., & Wager, T. D. (2011). Large-scale automated synthesis of human functional neuroimaging data. Nature Methods. doi:10.1038/nmeth.1635

Yellin, D., Berkovich-Ohana, A., & Malach, R. (2015). Coupling between pupil fluctuations and resting-state fmri uncovers a slow build-up of antagonistic responses in the human cortex. NeuroImage, 106, 414–427. Read. doi:10.1016/j.neuroimage.2014.11.034

Zacà, D., Hasson, U., Minati, L., & Jovicich, J. (2018). Method for retrospective estimation of natural head movement during structural mri. Journal of Magnetic Resonance Imaging, 48(4), 927–937.

Zhang, Y., Yan, A., Liu, B., Wan, Y., Zhao, Y., Liu, Y., … Liu, Z. (2018). Oculomotor performances are associated with motor and non-motor symptoms in parkinson’s disease. Frontiers in neurology, 9, 960.

